# Exercise induces Skeletal Muscle Methylome and Transcriptome changes, regardless of Age and COPD

**DOI:** 10.64898/2026.03.06.710054

**Authors:** Poojitha Rajasekar, Lorna Latimer, Linzy Houchen-Wolloff, Kamini Rakkar, Tim Constantin-Teodosiu, Julie L. MacIsaac, Lisa M. McEwen, Chen Xi Yang, Tillie-Louise Hackett, Bhavesh Popat, Despina Constantin, Michael S. Kobor, Michael C Steiner, Paul Greenhaff, Charlotte E Bolton, Rachel L Clifford

## Abstract

Skeletal muscle atrophy and deconditioning contribute to functional limitation and disability in COPD. While transcriptome and DNA methylation changes accompany exercise in healthy muscle, their interaction with COPD status and ageing, and integrative analyses of methylome–transcriptome responses have not been explored. We performed gene expression and DNA methylation profiling in skeletal muscle of sedentary volunteers with COPD, age-matched older adults, and younger healthy individuals, before and during (1,4 and 8 weeks) supervised aerobic exercise training and after four weeks of detraining. Exercise induced transcriptomic and DNA methylation changes, but these responses were unaffected by COPD status or age. Subsequent analysis focusing on temporal exercise effects independent of disease or age revealed differential transcriptomic changes across time points, a subset of which significantly associated with DNA methylome alterations. Transient transcriptomic changes not linked to DNA methylation were enriched for inflammatory and oxidative stress pathways, whereas persistent methylation-associated adaptations were related to immunomodulation and tissue remodelling. Together, this study provides insight into molecular mechanisms contributing to skeletal muscle adaptation to aerobic exercise training in sedentary individuals.

## Introduction

Physical activity well established to reduce the risk of chronic disease development (Marques et al, 2018) and progression of chronic illnesses is frequently associated with low levels of physical activity (Booth et al., 2012). Accordingly, aerobic exercise programs are an integral part of chronic lung disease treatment, because they enhance physiological capacity and resilience, thereby improving symptoms and quality of life (Pedersen & Saltin, 2015). Aerobic exercise induces both transient and long-term adaptive responses in skeletal muscle including molecular processes that modulate contractile protein composition (Cochran et al, 2014; Gonzalez et al, 2015), mitochondrial mass and function (Hood et al, 2015; Ringholm et al, 2023), intracellular signalling pathways (Benziane et al, 2008; Figueiredo et al, 2015; Hammond et al, 2019), substrate metabolism, and vascular function (Egan & Zierath, 2013).

Changes in gene expression in skeletal muscle are closely linked to functional adaptions during and following exercise training interventions (Brown, 2015; Contrepois *et al*, 2020; Egan & Zierath, 2013; Figueiredo *et al*, 2015; Perry *et al*, 2010). DNA methylation is a key regulatory mechanism that may underlie these transcriptional responses. DNA methylation involves the transfer of a methyl group to the C5 position of cytosine residues, forming 5-methylcytosine (Moore *et al*, 2013). This modification regulates gene expression in multiple chronic diseases (Brorson *et al*, 2022; Huang *et al*, 2014; Lalchungnunga *et al*, 2022; Morselli *et al*, 2022; Smyth *et al*, 2022), and has also been implicated in both acute and chronic responses to exercise (Hunter *et al*, 2019; Luttropp *et al*, 2013; Machado *et al*, 2021).

In skeletal muscle, targeted analysis has identified DNA methylation regulated gene expression alterations in several exercise-responsive genes, including peroxisome proliferator-activated receptor gamma coactivator 1-alpha gene *(PPARGC1A)* (Alibegovic *et al*, 2010), peroxisome proliferator-activated receptor gamma coactivator 1-alpha *(PGC-1α)*, pyruvate dehydrogenase kinase 4 *(PDK4)*, peroxisome proliferator-activated receptor gamma *(PPAR-δ)* (Barrès *et al*, 2012), thyroid adenoma associated gene *(THADA),* myocyte enhancer factor-2 *(MEF2A),* RUNT-related transcription factor 1 *(RUNX1),* and NADH dehydrogenase [ubiquinone] 1 subunit C2 *(NDUFC),* in response to 6 months of endurance training (Nitert *et al*, 2012). Additional work has identified DNA methylation changes in skeletal muscle function-related genes after both acute and chronic exercise training (Brown, 2015). Genome-wide DNA methylome and transcriptome profiling has further linked DNA methylation to gene expression changes associated with skeletal muscle atrophy (Laker *et al*, 2017) following acute (within 24 hours) and chronic (12 weeks) resistance exercise training. Notably, DNA methylation associated transcriptional changes can persist for at least seven weeks after exercise cessation, suggesting the presence of an “epigenetic memory” to resistance training (Seaborne *et al*, 2018; Turner *et al*, 2019). However, comprehensive longitudinal integration of DNA methylation and gene expression from matched skeletal muscle tissue samples across exercise and withdrawal timepoints remains limited, representing an important gap in understanding the molecular basis of skeletal muscle exercise adaption.

Chronic disease risk is strongly influenced by age, and DNA methylation patterns are highly age sensitive. Genome-wide DNA methylation profiling of skeletal muscle from young and older individuals has revealed widespread hypermethylation with ageing (Turner *et al*, 2020; Zykovich *et al*, 2014). DNA regions surrounding Homeobox (*HOX)* genes, key age-associated transcription factors, show reduced DNA methylation in aged skeletal muscle tissue and increased methylation variability in aged muscle-derived primary cells during differentiation compared to younger cells (Turner *et al*., 2020). More recently, development of a skeletal muscle epigenetic clock (Voisin *et al*, 2021) demonstrated that aerobic exercise training can shift the epigenetic and transcriptomic age of skeletal muscle towards a younger profile (Voisin *et al*, 2023). While these findings underscore an important role for DNA methylation in age-related skeletal muscle regulation, interpretation is complicated by the well documented age-associated decline in habitual physical activity and accompanying changes in skeletal muscle architecture and metabolism (Shur *et al*, 2021). Moreover, no study has systematically examined how age modifies DNA methylation-associated gene expression in skeletal muscle in the context of exercise adaption.

Chronic Obstructive Pulmonary Disease (COPD) is a debilitating lung disorder and a leading cause of global mortality (Soriano *et al*, 2020). Although primarily a pulmonary disease, COPD is associated with significant extra-pulmonary manifestations, including skeletal muscle dysfunction. Skeletal muscle dysfunction in COPD is characterised by reduced muscle strength, aerobic capacity and endurance and a shift from type I muscle fibres to type II (Gosker *et al*, 2007; Jarosch *et al*, 2016; Patel *et al*, 2014). Habitual physical activity declines with increasing disease severity (Watz *et al*, 2009) and greater sedentary time predicts mortality in COPD (Furlanetto *et al*, 2017). Ageing is also attributed to increased risk of developing COPD (Leon, 2011). Importantly, pulmonary rehabilitation incorporating supervised exercise improves endurance time, fatigue resistance and oxidative enzyme activity in COPD patients, leading to improved quality of life (Costes *et al*, 2015; Mador *et al*, 2014; Pothirat *et al*, 2015). Previous research has identified distinct mitochondrial functional and mass changes after eight weeks of aerobic exercise in healthy young individuals compared to healthy older individuals and COPD patients. However, no discernible age- or COPD-related differences were observed in exercise-induced expression of muscle fuel metabolism genes at moderate relative training intensities (60-70% peak work) (Latimer *et al*, 2022). Thus, despite strong evidence for the benefits of exercise in COPD rehabilitation, studies investigating DNA methylation-mediated gene regulation in response to aerobic training in this population are lacking. While altered DNA methylation has been reported in COPD in blood (Qiu *et al*, 2012; Yoo *et al*, 2015), peripheral blood mononuclear cells (Chen *et al*, 2021), whole lung tissue (Sundar *et al*, 2017), small airway epithelial cells (Vucic *et al*, 2014), and airway and parenchymal fibroblasts (Clifford *et al*, 2018), corresponding analyses in skeletal muscle have not been performed.

Here we performed retrospective analysis of vastus lateralis muscle tissue extracts obtained from an 8-week standardised, supervised aerobic exercise intervention (longitudinal timepoints: 1 week, 4 weeks and 8 weeks) followed by a 4-week exercise withdrawal period. Specifically, we generated matched transcriptome and DNA methylome profiles in vastus lateralis muscle samples of healthy young, COPD and age matched older individuals, to define genome-wide molecular adaptions to aerobic training, building on our previous work on mitochondrial function and fuel metabolism (Latimer *et al*., 2022). Here we have identified that age and COPD status did not significantly impact baseline or temporal transcriptional or DNA methylation responses to aerobic exercise or detraining. In contrast, exercise induced robust time-dependant changes in both the DNA methylome and transcriptome, with a subset persisting after four weeks of exercise withdrawal. Exercise-associated gene expression changes independent of DNA methylation were enriched for metabolic, oxidative stress response pathways, cytoskeletal, cell motility and adhesion pathways, whereas DNA methylation-associated transcriptional changes were predominantly enriched for immunomodulatory and muscle remodelling pathways across exercise weeks 1, 4 and 8. Together, these findings support a crucial role for DNA methylation in mediating exercise-induced transcriptional remodelling in skeletal muscle.

## Results

### Study cohort

The study cohort included three independent sedentary groups: 20 individuals with COPD, 10 physically in-active age-matched older healthy controls, and 9 physically in-active younger healthy controls (Table 1). Participants underwent eight weeks of supervised cycling-based aerobic exercise training (three 30 min sessions per week at a workload corresponding to 65% V′O2peak). Training intensity was reassessed at week 4 and workload adjusted if V′O2peak workload had increased. This exercise intervention was followed by four-week exercise withdrawal period. Metabolic and physiological endpoints measured throughout the study have been previously described (Latimer *et al*., 2022). Vastus lateralis skeletal muscle biopsies were collected in the fasted, resting state after 1, 4 and 8 weeks of exercise training and again after four weeks of exercise withdrawal for integrated DNA methylome and transcriptome profiling (Visual Synopsis). Patient demographics, statistical models, and covariates used for transcriptome and methylome analyses are detailed in Table 2 and 3.

**Visual Synopsis:**
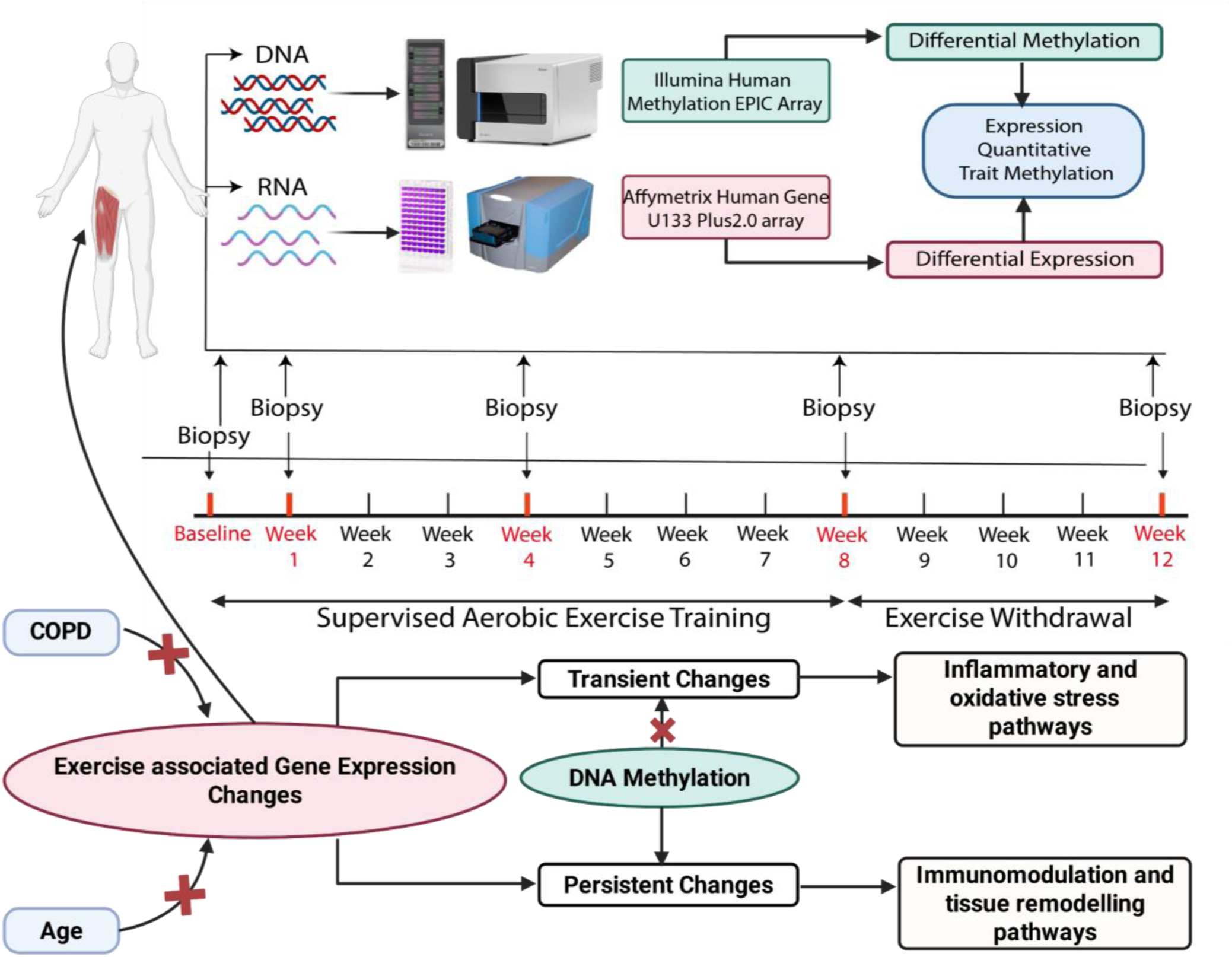
Supervised exercise training methylome and transcriptome study design. Skeletal muscle (Vastus Lateralis) microbiopsy was performed at resting and fasted state, 24 hours post exercise training. DNA and RNA extracted from the biopsy was subject to methylome and transcriptome profiling using Illumina Human Methylation EPIC array and Affymetrix Human Gene U133 Plus 2.0 array. The data was analysed to identify differential methylation and expression. The association between differential expression and methylation was investigated using expression quantitative trait methylation analysis. Transient transcriptomic changes not linked to DNA methylation were enriched for inflammatory and oxidative stress pathways, whereas persistent methylation-associated adaptations were related to immunomodulation and tissue remodelling.

**Table 1:** Study cohort demographics. (mean ± SD). BMI – Body Mass Index; FEV_1_ - Forced Expiratory Volume; FVC – Forced Vital Capacity; RV – Residual Volume; TLC – Total Lung Capacity; TLCO - Transfer Factor of the Lung for Carbon Monoxide; FFMI – Fat Free Mass Index; VO_2_ – Oxygen Consumption.

| **Demographics** | **Young Healthy**<br>**(n=9)** | **Older**<br>**Healthy**<br>**(n=10)** | **COPD**<br>**(n=20)** |
| --- | --- | --- | --- |
| **Age, Years** | 28.0±5.1 | 70.7±5.1 | 70.2±5.9 |
| **Female, number** | 6 | 5 | 14 |
| **BMI, kg m<sup>-2</sup>** | 26±7.6 | 28.5±3.3 | 29.0±6.4 |
| **Smoking, Pack Years** | 0±0 | 0±8 | 40±24 |
| **FEV<sub>1</sub>, % predicted** | 112.6±20.6 | 98.3±9.2 | 55.6±16.2 |
| **FEV<sub>1</sub>/FVC, ratio** | 81.61±9.32 | 74.5±4.4 | 44.6±12.1 |
| **RV, & predicted** | 107.9±48.9 | 98.4±19.2 | 157.3±44.9 |
| **TLC, % predicted** | 107.6±15.0 | 104.3±11.9 | 123.1±19.7 |
| **RV/TLC, ratio** | 26.8±9.7 | 38.4±4.2 | 53.5±9.4 |
| **TLCO, % predicted** | 95.6±13.9 | 98.3±12.7 | 61.5±18.7 |
| **FFMI, kg m<sup>2</sup>** | 16.5±2.8 | 18.1±1.5 | 17.2±2.7 |
| **Isokinetic Strength, Nm** | 120.2±51.8 | 106.5±32.9 | 76.8±29.0 |
| **Values at baseline incremental CPET testing at peak VO<sub>2</sub>** |  |  |  |
| **Workload, Watts** | 167.0±43.6 | 113.0±26.4 | 71.1±32.5 |
| **VO<sub>2</sub>, ml lean mass<sup>-1</sup> min<sup>-1</sup>** | 44.7±5.6 | 29.7±4.0 | 24.0±7.4 |

**Table 2:** Table of data analyses models used in differential DNA methylation analysis, specifying the purpose of analyses, samples used, and details of statistical tests performed on key demographic features to identify potential covariates for the model.

| DNA Methylation Analysis | Samples | Number of Samples |  |  |  |  | Significant Difference in Demographics | Linear Model Design |
| --- | --- | --- | --- | --- | --- | --- | --- | --- |
|  |  | Baseline | Exercise Week 1 | Exercise Week 4 | Exercise Week 8 | Exercise Withdrawal |  |  |
| Effect of COPD at Baseline | COPD (n=20) | 20 | - | - | - | - | Smoking Status (p=0.0003537) | ~Disease+ Smoke |
|  | Old Healthy (n=10) | 10 | - | - | - | - |  |  |
| Effect of Age at Baseline | Old Healthy (n=10) | 10 | - | - | - | - | Age (p=6.56e-14) | ~Age |
|  | Young Healthy (n=9) | 9 | - | - | - | - |  |  |
| Interaction effect of COPD and Exercise at 5 time points | COPD (n=94) | 20 | 17 | 19 | 20 | 18 | Smoking Status (p=0.0003537) | ~Exercise* Disease +Smoke + Age |
| Interaction effect of Age and Exercise at 5 time points | Old healthy (n=50) | 10 | 10 | 10 | 10 | 10 | Age (p=6.56e-14) | ~Exercise* Age +Disease +Smoke |
| Effect of exercise independent of COPD and age at 5 time points | Young Healthy (n=47) | 9 | 9 | 9 | 10 | 10 | Smoking Status (p=0.0003537); Age (p=6.56e-14) | ~Exercise+ Disease+ Age+Smoke |

**Table 3:** Table of data analyses models used in differential gene expression analysis, specifying the purpose of analyses, samples used and details of statistical tests performed on key demographic features to identify potential covariates for the model.

| Gene Expression Analysis | Samples | Number of Samples |  |  |  |  | Significant Difference in Demographics | Linear Model Design |
| --- | --- | --- | --- | --- | --- | --- | --- | --- |
|  |  | Baseline | Exercise Week 1 | Exercise Week 4 | Exercise Week 8 | Exercise Withdrawal |  |  |
| Effect of COPD at Baseline | COPD (n=19) | 19 | - | - | - | - | Smoking Status (p= 0.0001255) | ~Disease+ Smoke |
|  | Old Healthy (n=10) | 10 | - | - | - | - |  |  |
| Effect of Age at Baseline | Old Healthy (n=10) | 10 | - | - | - | - | Age (p=6.56e-14) | ~Age |
|  | Young Healthy (n=9) | 10 | - | - | - | - |  |  |
| Interaction effect of COPD and Exercise at 5 time points | COPD (n=74) | 19 | 19 | 19 | 19 | 17 | Smoking Status (p=0.0001255) | ~Exercise* Disease +Smoke + Age |
| Interaction effect of Age and Exercise at 5 time points | Old healthy (n=49) | 10 | 9 | 10 | 10 | 10 | Age (p=6.56e-14) | ~Exercise* Age +Disease +Smoke |
| Effect of exercise independent of COPD and age at 5 time points | Young Healthy (n=45) | 10 | 7 | 9 | 9 | 10 | Smoking Status (p=0.0001255); Age (p=6.56e-14) | ~Exercise+ Disease+ Age+Smoke |

### DNA methylation and gene expression are not associated with COPD status or age at baseline

To determine whether baseline skeletal muscle DNA methylation or gene expression differed by COPD status or age, global DNA methylation and genome-wide gene expression was assessed using linear regression analysis. No significant differences in DNA methylation or gene expression were observed between individuals with COPD and age-matched older healthy controls (BH FDR<9x10^-8^; COPD n=20, older healthy individuals n=10), suggesting no fundamental baseline differences in skeletal muscle molecular profiles associated with COPD. Similarly, no significant age-associated differences in DNA methylation or gene expression were detected between younger and older healthy controls (BH FDR<9x10^-8^; Younger Healthy n=9; Older Healthy n=10).

### COPD status and age have no impact on exercise associated differences in gene expression or DNA methylation

We next examined whether COPD status or age modified exercise-induced molecular responses using interaction term linear regression models. No significant exercise x COPD status or age on exercise were detected for either DNA methylation or gene expression interactions (BH FDR<9x10^-8^) at any timepoint. Together, these analyses suggest that exercise-associated molecular adaptions occur largely independently of COPD status and age.

### DNA methylation and gene expression changes are associated with exercise and its withdrawal

Having established no detectable impact of COPD status or age on DNA methylation or gene expression changes exercise responses, we next investigated exercise-associated DNA methylation and gene expression changes over time. All samples were included in the analysis, with COPD status and age as covariates.

A total of 108,983, 49,955, 168,092 and 99,561 CpGs were identified as differentially methylated positions (DMPs) relative to baseline, at 1 week (Fig 1A), 4 weeks (Fig 1B), 8 weeks of exercise (Fig 1C) and 4 weeks post-exercise withdrawal (Fig 1D), respectively (BH FDR p<9x10^-8^) (Appendix Table S1). Increased DNA methylation predominant at week 1, whereas effect direction was more balanced at later time points (Appendix Table S1). The magnitude of change in DNA methylation beta values ranged from -0.21 to 0.21 at week 1, -0.23 to 0.19 at week 4, -0.28 to 0.28 at week 8 and -0.27 to 0.21 post-exercise withdrawal (Fig 1A-D).

**Figure 1:**
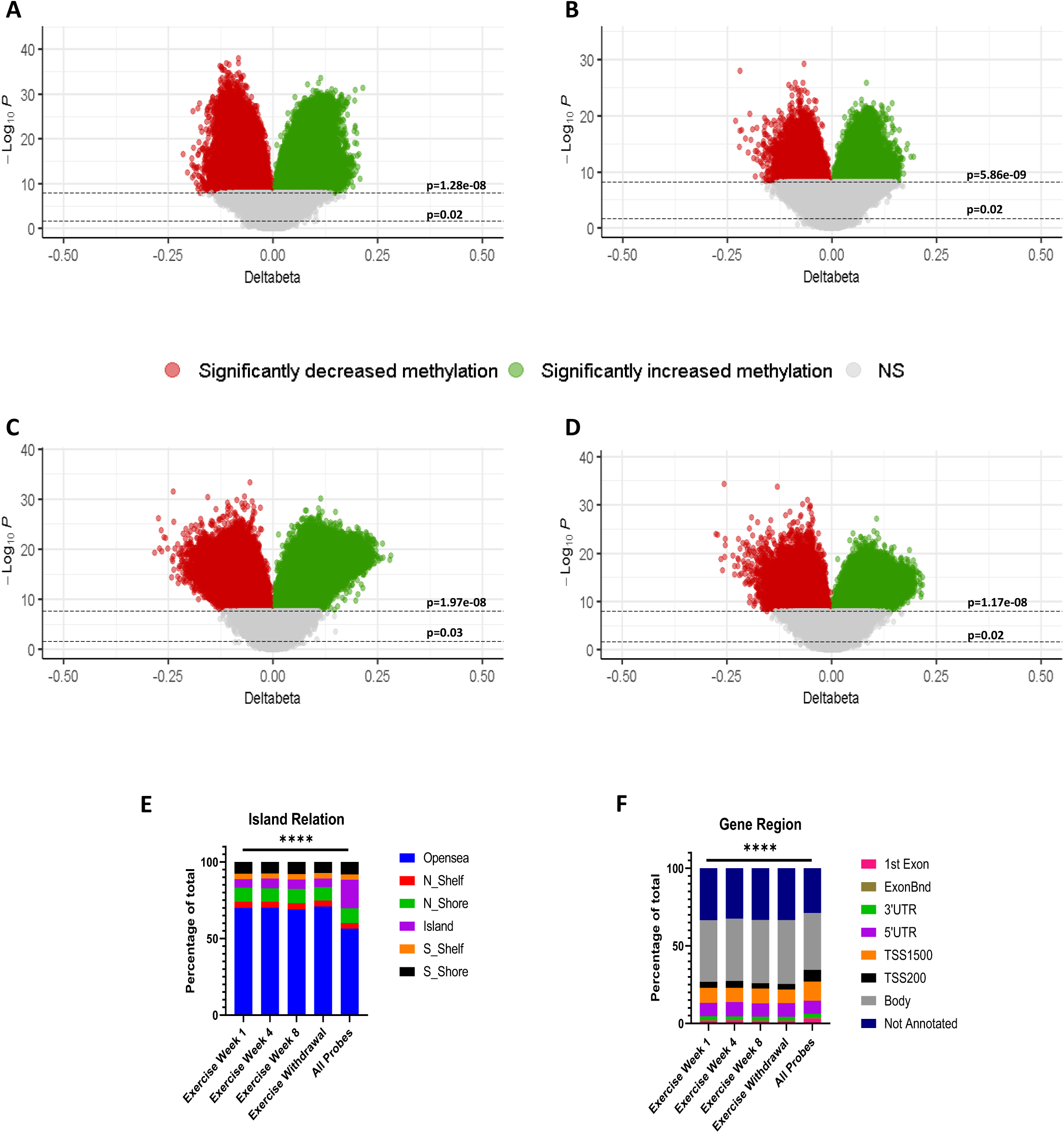
Effect of exercise on skeletal muscle DNA methylation. Plot of 766569 DNA methylation probes used in the analysis of response to exercise at **[A]** week 1, **[B]** week 4, **[C]** week 8 and **[D]** 4 weeks of exercise withdrawal. Red/Green = Benjamini-Hochberg FDR < 9e-8. The horizontal dotted lines indicate the p value corresponding to Benjamini-Hochberg FDR < 0.05 (bottom) and 9e-8(top). Red = decreased methylation at exercise timepoint relative to baseline, green = increased methylation at exercise timepoint relative to baseline. **[E]** Contingency plot depicting distribution of CpG Island locations and **[F]** gene region locations of differentially methylated positions. Chi-Square test demonstrated significant (**** = p<0.0001) difference in distribution across the timepoints in comparison to all probes used in analysis.

At all time points, DMPs were significantly enriched in open sea regions (70%; Fig 1E, Chi-Square test, p<0.0001) and gene bodies (40%; Fig 1F, Chi-Square test, p<0.0001), relative to the distribution of all analysed probes. Open sea regions are isolated CpGs sites in the genome shown to display an inverse relationship with gene expression (Dayeh *et al*, 2014) whereas gene body DNA methylation is frequently positively correlated with gene expression and linked to chromatin accessibility (Jjingo *et al*, 2012).

A total of 39,112 CpGs were differentially methylated across all timepoints (Fig 2A) with highconsistent directionality (Fig EV1A, Pearson Correlation ≥0.97; p<0.0001). At week 4, 88% (n=44,002) of DMPs overlapped with week 1 (Fig 2A). At week 8, 50% (n=85,412) and 29% (n=48,327) of DMPs overlapped with weeks 1 and 4, respectively (Fig 2A), with increasing concordance in effect size over time (Fig 2B, Pearson correlation: Exercise week 1 = 0.97 and Exercise week 4=0.99). Following exercise withdrawal, 64% (n=63,922), 44% (n=44,230), and 97% (n=96,199) of DMPs overlapped with weeks 1, 4 and 8 of exercise, respectively again with strengthening correlations (Fig 2A, Pearson correlation: Exercise week 1 = 0.97, Exercise week 4=0.991 and Exercise week 8=0.996) in magnitude over time (Fig 2C). These data indicate a rapid DNA methylation response within one week of exercise that progressively stabilizes with continued training.

**Figure 2:**
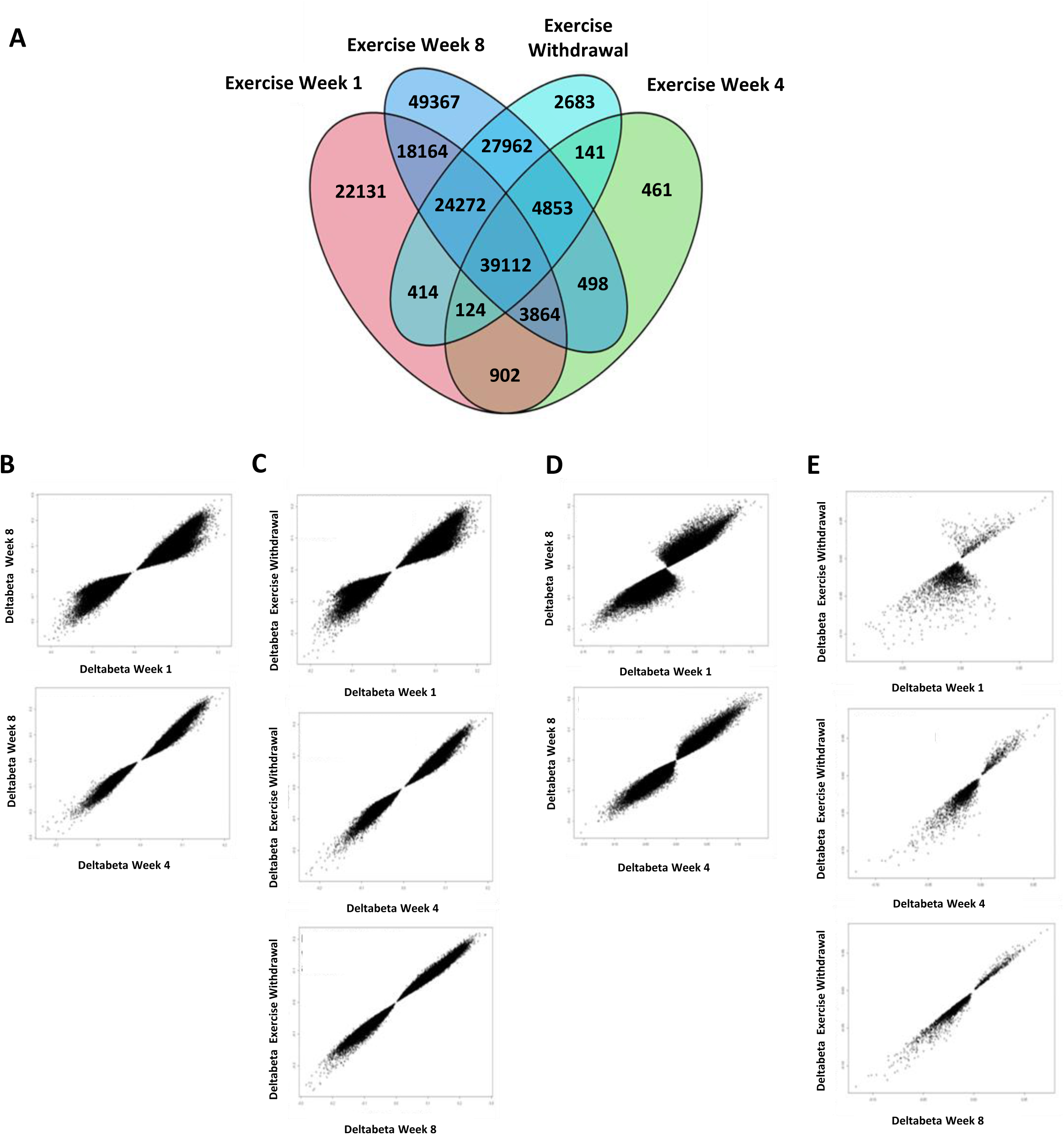
Comparing effect size, overlap and direction of change between exercise associated skeletal muscle DNA methylation changes across time points. **[A]** Venn diagrams depicting overlap between significant differentially methylated positions (Benjamini-Hochberg FDR < 9e-8) across time points**. [B-E]** Correlation plots depicting profiles of **[B]** week 8 hits that were present at weeks 1 and 4, **[C]** exercise withdrawal hits that were present at weeks 1,4 and 8, **[D]** new week 8 hits at weeks 1 and 4, **[E]** new exercise withdrawal hits at weeks 1, 4 and 8.

Notably, 46% (n=77,329) and 3% (n=2,683) (Fig 2A) of DMPs at week 8- and post-exercise withdrawal, respectively, were newly identified. Correlations of these CpGs acress earlier time points suggested gradual CpG methylation shifts that reached statistical significance later in the intervention and post-exercise (Fig 2D, Pearson correlation: Exercise week 1 = 0.94 and Exercise week 4=0.98; Fig 2E, Pearson correlation: Exercise week 1 = 0.71, Exercise week 4=0.95 and Exercise week 8=0.98).

#### Exercise Induces sustained transcriptional remodelling

We next assessed exercise-associated gene expression changes. A total of 4413, 2081, 1682 and 4406 probes were differentially expressed relative to baseline at week 1 (Fig 3A), week 4 (Fig 3B), week 8 (Fig 3C), and post-exercise withdrawal (Fig 3D), respectively (BH FDR p<9x10^-8^), corresponding to 2570, 1271, 1031 and 2549 unique annotated genes (Appendix Table S2). Gene expression changes were predominantly upregulated in response to exercise at all time points relative to baseline (Fig 3A-D and Appendix Table S2).

**Figure 3:**
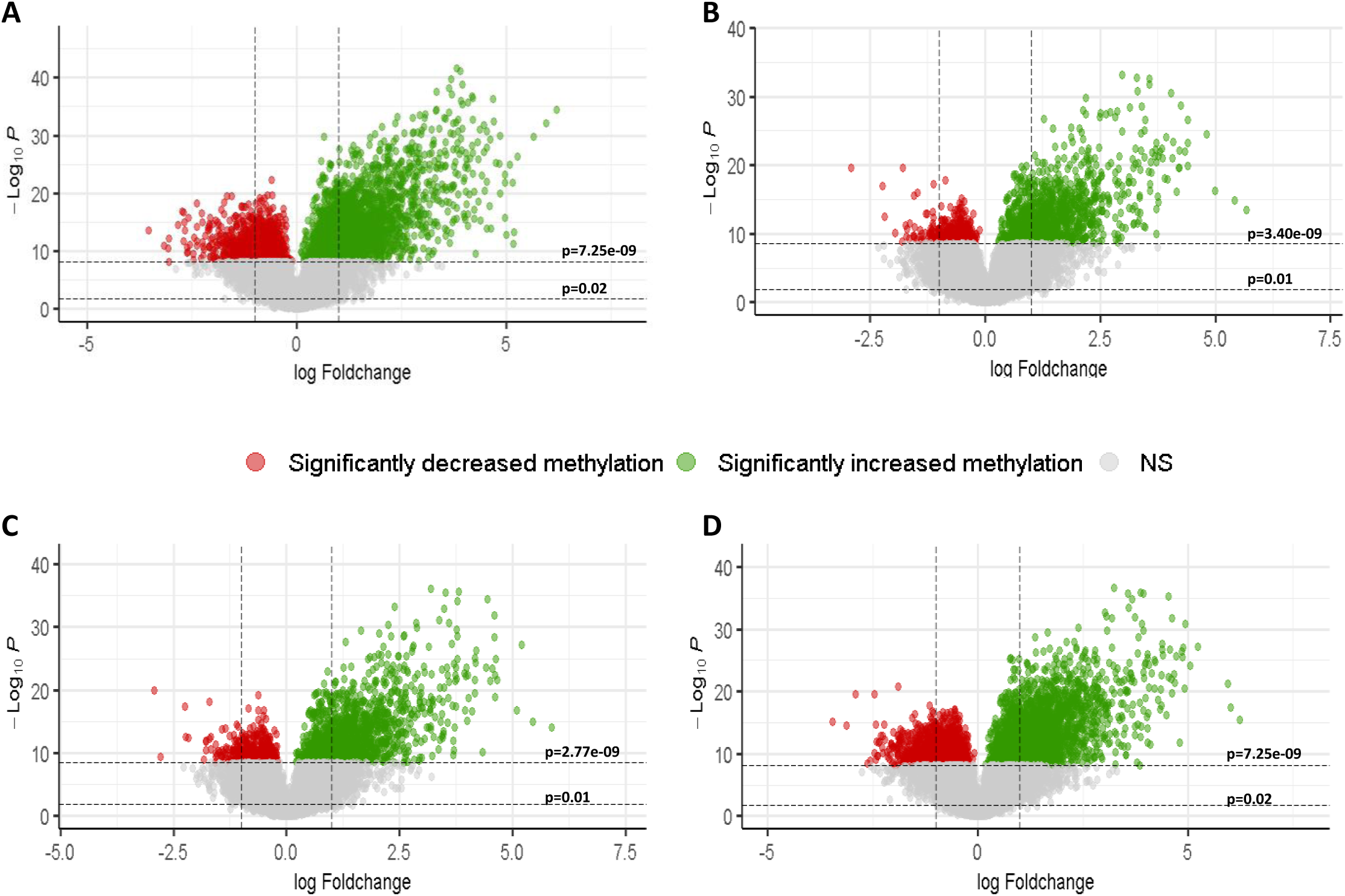
Effect of exercise on skeletal muscle gene expression. **P**lot of 54,675 transcript probes used in the analysis of response to exercise at **[A]** week 1, **[B]** week 4, **[C]** week 8 and **[D]** 4 weeks of exercise withdrawal. Red/Green = Benjamini-Hochberg FDR < 9e-8. The horizontal dotted lines indicate the p value corresponding to Benjamini-Hochberg FDR < 0.05 (bottom) and 9e-8(top). Red = decreased expression at exercise timepoint relative to baseline, green = increased expression at exercise timepoint relative to baseline.

1244 probes were differentially expressed across all timepoints (Fig 4A), with a consistent direction of change, indicative of a sustained transcriptomic response to exercise, that builds over time and remains upon cessation of exercise (Fig EV1 B). At week 8, 80% (n=1349) and 88% (n=1488) of significantly differentially expressed genes (DEGs) were present at weeks 1 and 4 respectively (Fig 4A), with increased correlation over time (Fig 4B, Pearson correlation: Exercise week 1 = 0.95 and Exercise week 4=0.99). At 4 weeks post exercise withdrawal 61% (n=2675), 44% (n=1946) and 38% (n=1618) (Fig 4A) of significantly DEGs were present at weeks 1, 4 and 8 of exercise respectively (Fig 4C, Pearson correlation: Exercise week 1 = 0.97, Exercise week 4=0.99 and Exercise week 8=0.99). Similar to the trend observed in DNA methylation, there is an initial response at exercise week 1 that progresses to become more stable over time with some changes remaining despite exercise withdrawal.

**Figure 4:**
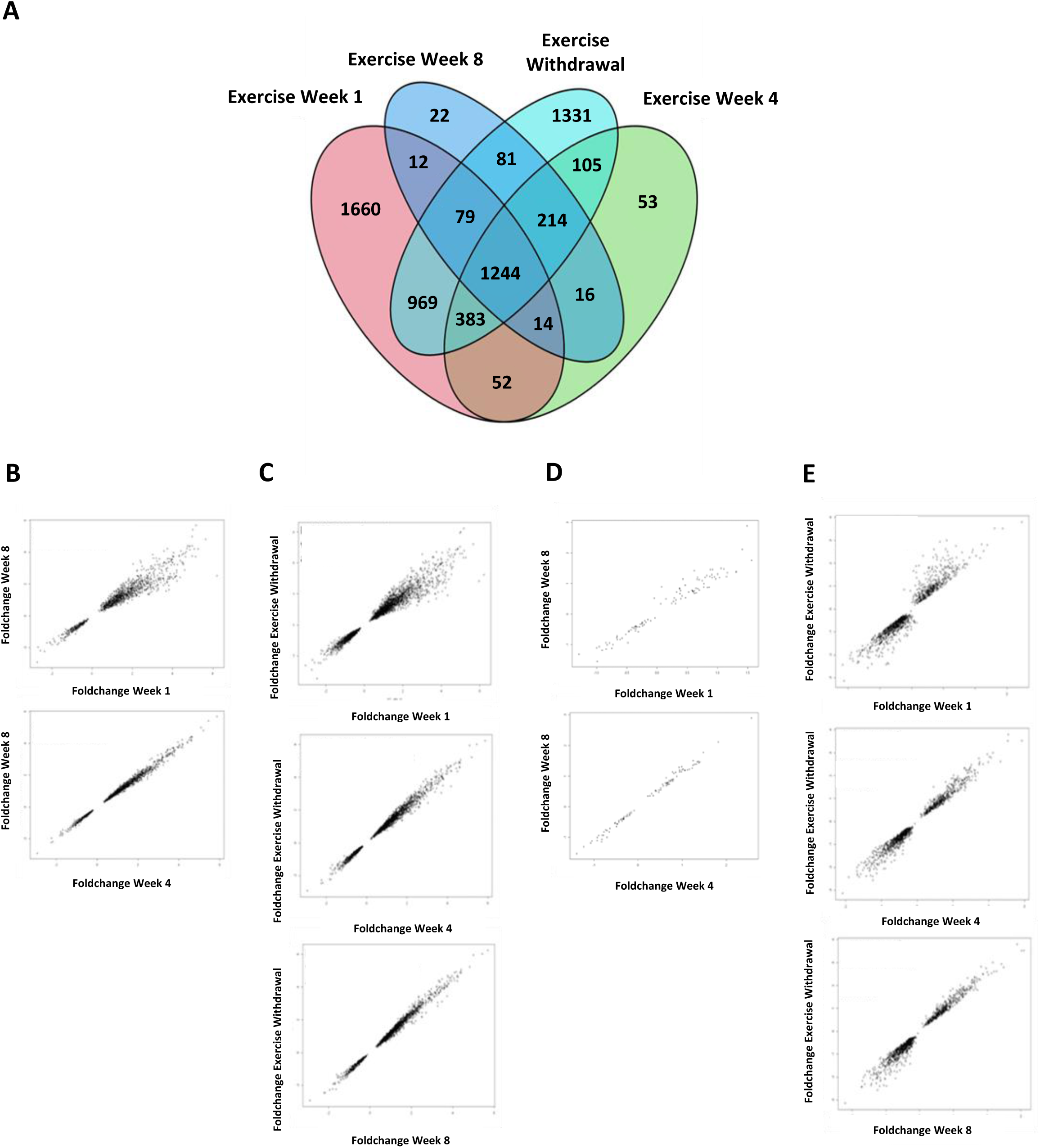
Comparing effect size, overlap and direction of change between exercise associated skeletal muscle gene expression changes across time points. **[A]** Venn diagrams depicting overlap between significant DEGs (Benjamini-Hochberg FDR < 9e-8) across time points**. [B-E]** Correlation plots depicting profiles of **[B]** week 8 hits that were present at weeks 1 and 4, **[C]** exercise withdrawal hits that were present at weeks 1,4 and 8, **[D]** new week 8 hits at weeks 1 and 4, **[E]** new exercise withdrawal hits at weeks 1, 4 and 8.

At 8- and 4-weeks post exercise withdrawal, 6% (n=103) and 30% (n=1331) (Fig 4A) of the DEGs were newly identified. Expression of the newly identified genes at exercise week 8 and exercise withdrawal increased in correlation over time, indicative of a gradual response to exercise in these genes, reaching significance at the later time point and upon exercise withdrawal (Fig 4D, Pearson correlation: Exercise week 1 = 0.97 and Exercise week 4=0.99; Fig 4E, Pearson correlation: Exercise week 1 = 0.95, Exercise week 4=0.98 and Exercise week 8=0.98).

Pathway analysis provided insight into the functional capacity of exercise-associated gene expression changes. Exercise responsive genes enriched to 151, 110, 89 and 115 significant pathways (p<0.05 and absolute(z-score)>1) for exercise weeks 1, 4, 8 and exercise withdrawal respectively (Appendix Table S3). 43 pathways were exclusive to exercise week 1 (Fig 5A) and predominantly comprised activation of pro-inflammatory pathways (macrophage classical activation signalling pathway, *IL-2* and *IL-13* signalling, inflammasome, interferon signalling and production of nitric oxide and reactive oxygen species in macrophages), metabolism and oxidative stress response pathways (sirtuin signalling, Glutathione redox reactions II, Hypoxia-inducible factor 1-alpha (*HIF-1α*) signalling) and muscle growth and tissue remodelling pathways (Insulin-like growth factor (*IGF1*) signalling, Eukaryotic Initiation Factor 2 (*EIF2*) Signalling) (Fig 5B, square points) suggesting an initial acute and transient activation of inflammatory, metabolic, oxidative stress and muscle activation pathways in response to exercise. There were only seven and two pathways that enriched exclusively for week 4 and 8 (Fig 5A), with a general transition to pathways that enriched across all timepoints. Specifically, 65 pathways were consistent across all time points (Fig 5A). Generally, these consistent pathways displayed higher significance and a greater z-score (Fig 5B, diamond points) than time-point specific pathways indicating early onset and long-lasting effects of exercise. They predominantly comprised immune and tissue remodelling pathways. Immune pathways included activation of some anti-inflammatory pathways (*IL-4* signalling, *Th2* and *IL-4* mediated macrophage alternative activation signalling pathway,*CXCR4* signalling) and pro-inflammatory responses (activation of *IL-8* signalling, *Th1* pathway, T cell receptor signalling, leukocyte extravasation signalling and inhibition of Cytotoxic T Lymphocyte antigen 4 (*CTLA4*) signalling and the *IL-10* pathway) (Fig 5B, green diamond points). Tissue remodelling pathways included cytoskeleton and cell adhesion modulating pathways including integrin signalling, actin cytoskeleton signalling, remodelling of epithelial adherens junction and Integrin-linked kinase (*ILK*) signalling (Fig 5B, purple and blue diamonds) potentially linked to increasing muscle mass and strength (Boppart & Mahmassani, 2019). Furthermore, inhibition of Rho GDP-dissociation inhibitor (*RHOGDI*) signalling and activation of Ras-related C3 botulinum toxin substrate (*RAC*), (Fig 5B) may promote glucose uptake (Moller *et al*, 2023), muscle fibre adaptation and immune cell migration to tissue repair (Kann *et al*, 2022), essential for training responsiveness and recovery. Activation of fibrosis tissue repair and remodelling signalling pathways (apelin liver signalling, Glycoprotein VI (*GP6*) signalling, pulmonary healing signalling, pulmonary fibrosis idiopathic signalling and hepatic fibrosis signalling) (Fig 5B, purple diamond) was also observed, indicative of adaptive tissue reorganisation and repair in response to exercise induced metabolic demand and oxidative stress that continues after exercise cessation.

**Figure 5:**
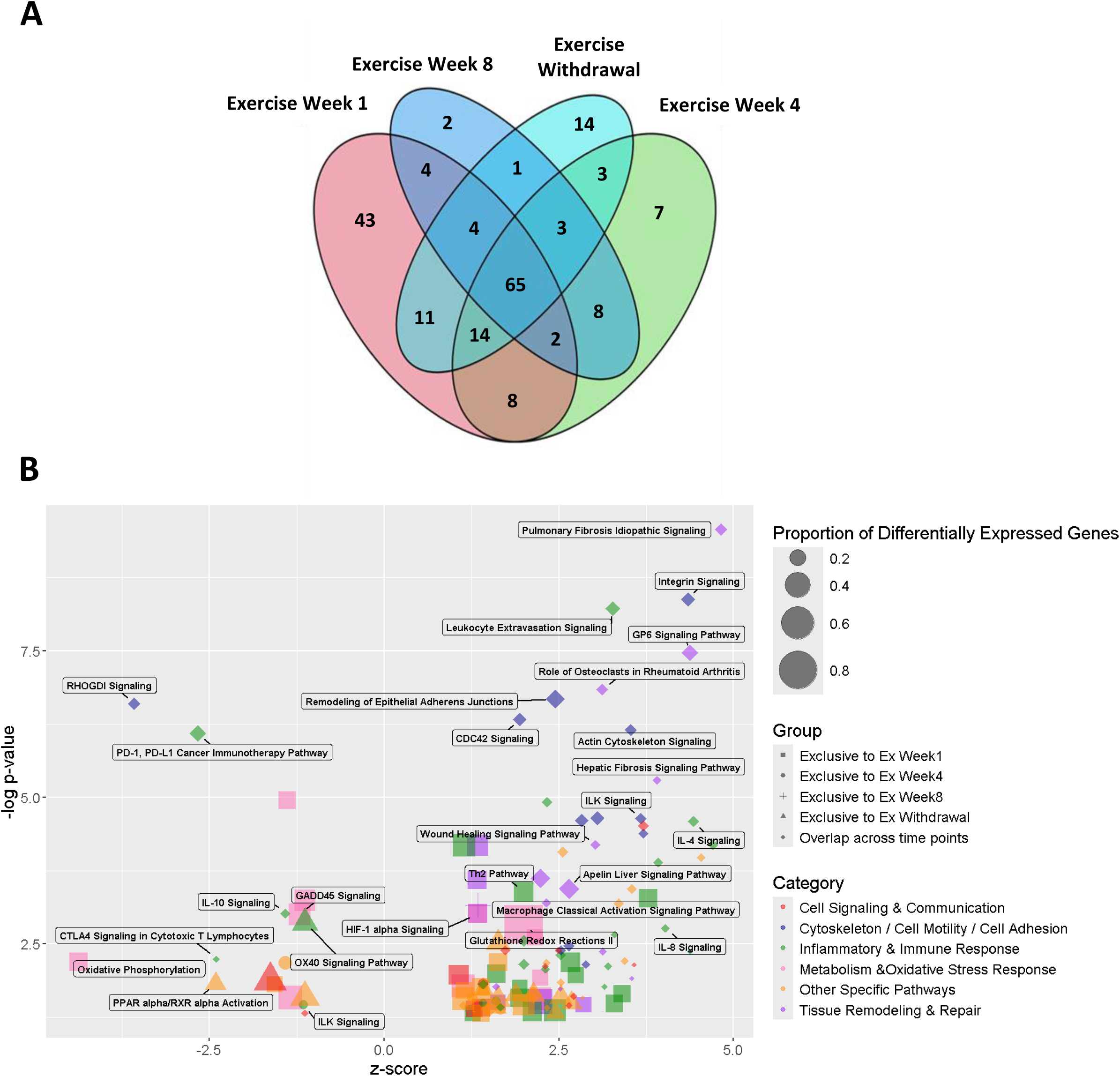
Pathway analysis of differentially expressed genes performed using Qiagen Ingenuity Pathway Analysis software. **[A]** Venn diagram depicting overlap of pathways enriched to DEGs at each time point. **[B]** A bubble plot of functional pathway associations that were exclusively enriched at each time point and those that were enriched across all time points, depicted by different shapes. The plot depicts −log(P value) and z-score on the axes, with the size of the bubble indicating the proportion of DEGs in the dataset comprised in the pathway.

Overall, pathway analysis of DEGs in response to exercise revealed an initial transient pro-inflammatory response and tissue remodelling activation that progressively shifted to an enrichment for a balanced pro and anti-inflammatory response, anti-fibrotic, tissue repair, myogenic and myokine response by 8 weeks of exercise. There was consistent activation of myogenesis mediating pathways throughout the exercise time course that was sustained even after exercise withdrawal.

Upstream regulator analysis was conducted to identify the key regulatory molecules and pathways that may be activated or inhibited to drive the observed changes in gene expression. 195, 199, 186 and 191 significant upstream regulators (p<0.05) were enriched at exercise weeks 1, 4, 8 and exercise withdrawal respectively (Appendix Table S5). 32 upstream regulators were transiently associated to one week of exercise (Fig 6A) and included activation of interferon regulatory factor 7 (*IRF7*) and inhibition of PR/SET domain 1 (*PRDM1*) (Fig 6B) associated with pro-inflammatory interferon, *IL-13* and *IL-2* signalling pathways that were transiently activated at exercise week 1 (Fig 5B) (Akman et al, 2021; Ma et al, 2023). In addition, histone deacetylases 1 and 5 (*HDAC1* and *HDAC5*), important epigenetic modulators of gene expression, were also enriched exclusively to exercise week 1. Regulation of DEGs exclusive to exercise week 4 enriched to 13 upstream regulators (Fig 6A) including Nuclear hormone receptor 4A (*NR4A1*) which regulates glucose metabolism (Fu et al, 2007) and skeletal muscle growth (Tontonoz et al, 2015) and is upregulated in response to exercise (Kanzleiter et al, 2009), Neurogenic locus notch homolog protein 2 (*NOTCH2*) and transcription factor 7-like 1 (*TCF L1*) which have known roles in myogenesis (Fujimaki et al, 2018) (Fig 6B, green circle), suggesting the onset of tissue remodelling signals. 17 upstream regulators (Fig 6A) were enriched only at exercise week 8 including more myogenesis associated transcription factors: transcription factor 4 (*TCF4*) crucial for muscle fibre type regulation, (Mathew et al, 2011), Recombination Signal Binding Protein for Kappa J (*RBPJ*) a *NOTCH* mediator essential for maintenance of muscle progenitor cells (Vasyutina et al, 2007) and transcription factor 12 (*TCF12*) that works in synergy with Myogenic Differentiation (*MYOD*) to modulate chromatin architecture to enable skeletal muscle regeneration (Wang et al, 2022) (Fig 6B, green plus). This emphasised the ongoing muscle repair and remodelling through the course of exercise. Four weeks after detraining, regulation of DEGs enriched to 23 new upstream regulators (Fig 6A) including Paired Box 3 (*Pax3*) which is characteristic of satellite cells and regulates myogenic differentiation via MYOD (Buckingham, 2007), peroxisome proliferator-activated receptor (*PPARG*) associated with fibre type switching in response to exercise and regulation of lipid metabolism and inflammation in skeletal muscle (Crossland et al, 2021; Phua et al, 2018), and Parkin (*PRKN*), a ubiquitin ligase implicated in regulating mitochondrial function following exercise induced oxidative stress phenotype (Chen et al, 2018) (Fig 6B, triangle). This strengthens the observation of muscle adaptation and regeneration pathway activation post exercise withdrawal and reiterates the potential molecular memory of exercise that is sustained even after exercise cessation.

**Figure 6:**
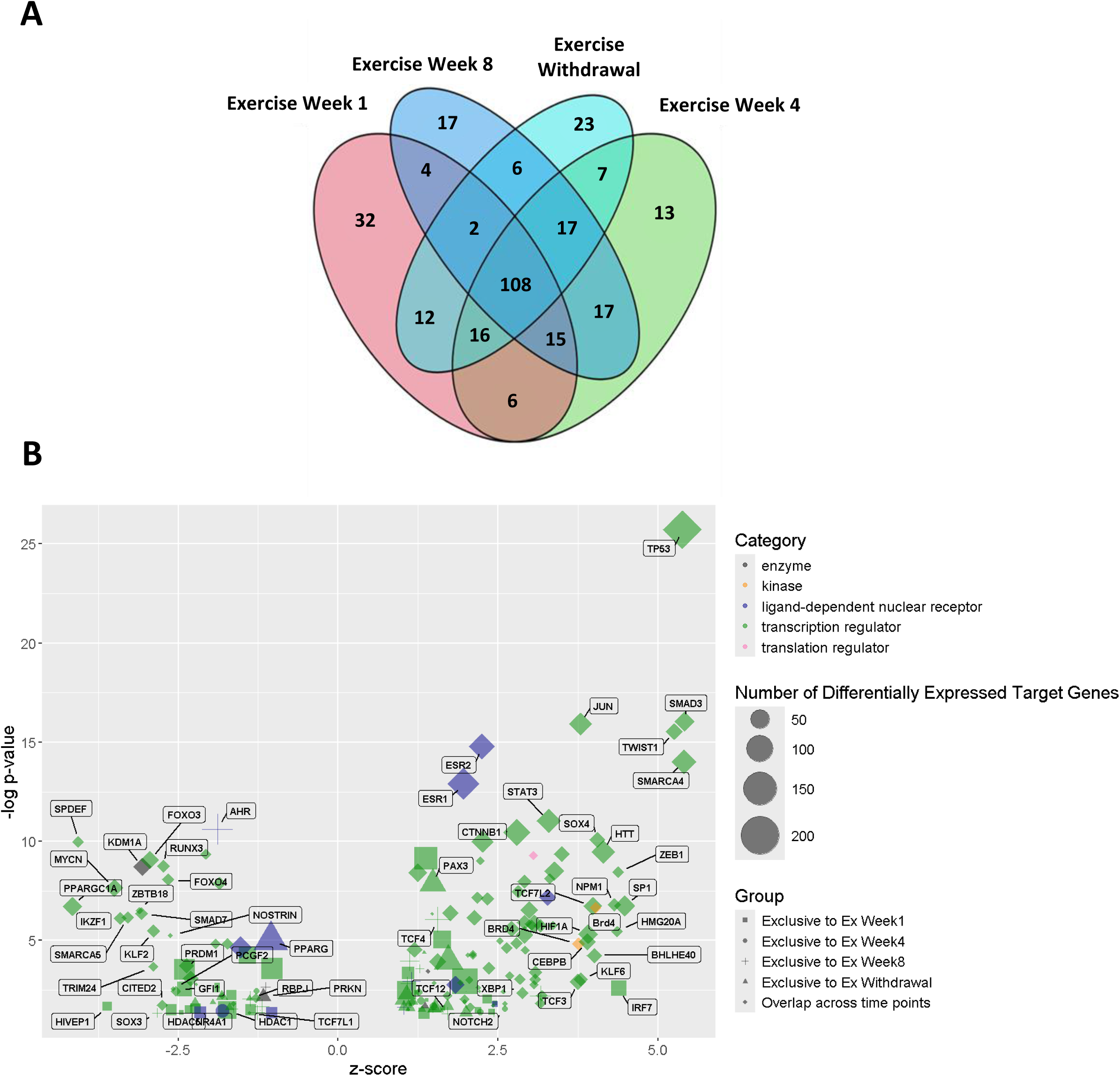
Upstream regulator enrichment analysis of differentially expressed genes performed using Qiagen Ingenuity Pathway Analysis software. **[A]** Venn diagram depicting overlap of upstream regulators enriched to DEGs at each time point. **[B]** A bubble plot of upstream regulator associations that were exclusively enriched at each time point and those that were enriched across all time points, depicted by different shapes. The plot depicts −log(P value) and z-score on the axes, with the size of the bubble indicating number of differentially expressed target genes in the dataset.

However, the most significantly enriched upstream regulators were those that spanned all time points. Specifically, 108 upstream regulators (Fig 6A) predominantly comprised of transcription regulators including, tumour protein p53 (*TP53*), associated with exercise capacity and aerobic metabolism (Park et al, 2009; Wang et al, 2012); myelocytomatosis oncogene (*MYC*), linked to age and exercise associated skeletal muscle remodelling (Jones et al, 2023), and SWI/SNF-related, matrix-associated, actin-dependent regulator of chromatin, subfamily A, member 4 (*SMARCA4*), a chromatin remodelling gene involved in early transcriptional remodelling of skeletal muscle (Terry et al, 2018) (Fig 6B, green diamond). Endoplasmic reticulum oxidative stress regulating transcription factor, X-box binding protein 1 (*XBP1*) (Alasiri et al, 2018) and mitochondrial respiration and exercise linked angiogenesis inducing Forkhead box O3 (*FOXO3*) (Sanchez, 2015) were enriched most significantly at exercise week 1 and remained significant, across time points (Fig 6B, green diamond). This observation links with the hypoxia and stress response pathways exclusively enriched to exercise week 1 including HIF-1α signalling, Growth Arrest and DNA damage-inducible 45 (*GADD45*) signalling and the Endoplasmic Reticulum stress pathway, implicating an oxidative stress response triggered in early exercise. Subsequently, TWIST family bHLH transcription factor 1 (*TWIST1*), Signal Transducer and Activator of Transcription 3 (*STAT3*) and SMAD family member 3 (*SMAD3*), associated with epithelial mesenchyme transition (Wang et al, 2017) were significant at week 1 but increasingly more significant from exercise week 4 and persisted after exercise withdrawal reiterating a tissue remodelling response to exercise.

### Exercise associated differential gene expression and DNA methylation are associated

Having identified exercise associated DMPs and DEGs we were interested in identifying any association between CpG methylation and gene expression via expression quantitative trait methylation (eQTM) analysis using DMPs and DEGs at each time point independently (Table 4).

**Table 4:**
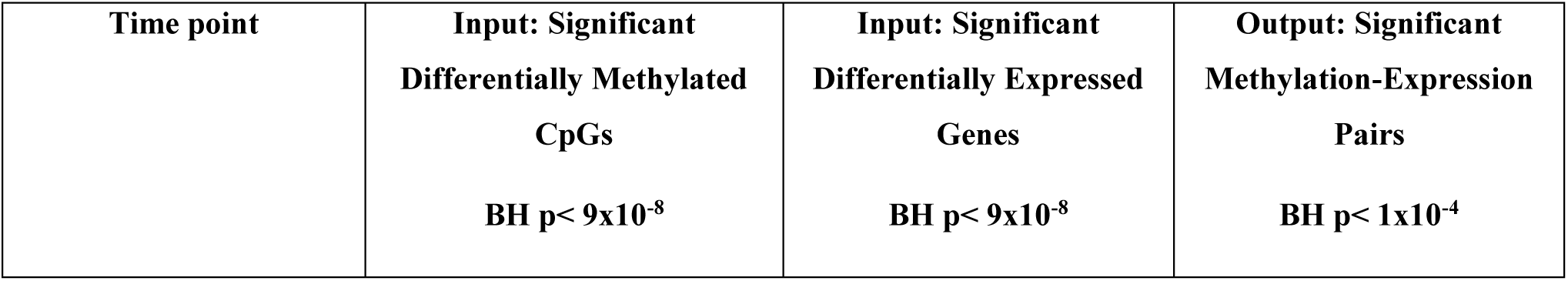
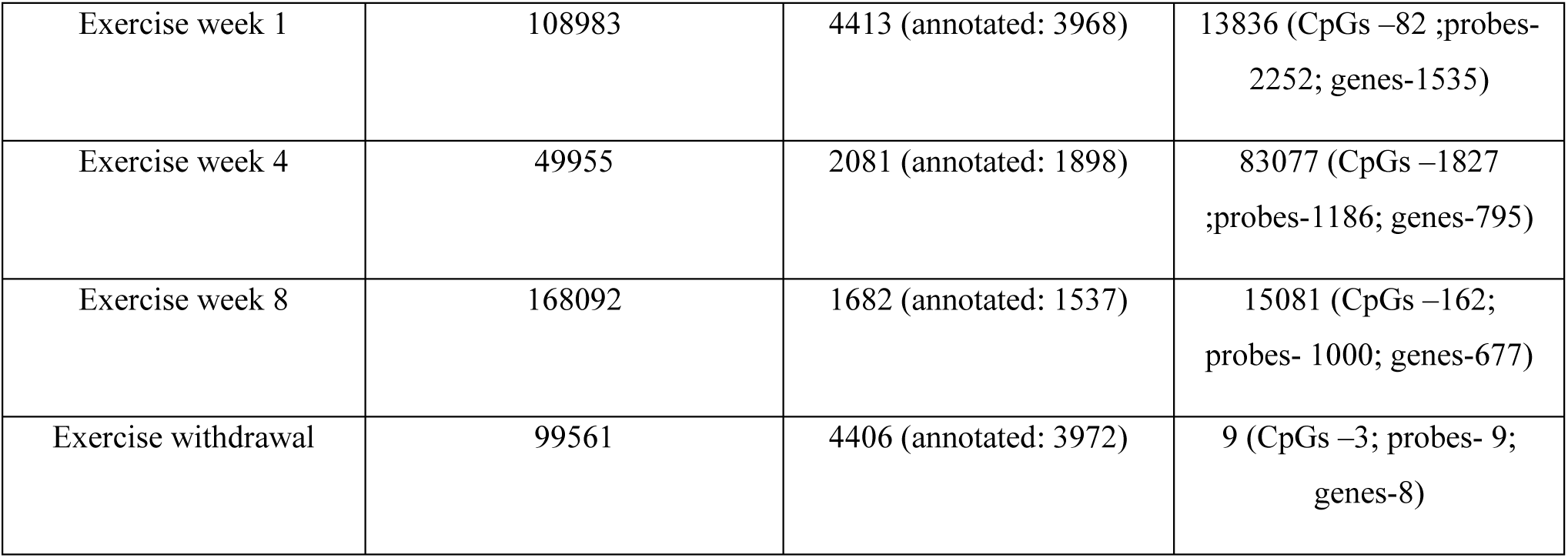
Expression Quantitative Trait Methylation (eQTM) input and results. The table depicts the number of significant differentially methylated CpGs and DEGs given as input into the eQTM model and the resulting number of significant (Benjamini-Hochberg FDR < 1e-4) methylation-expression pairs identified at each time point. The output column details the number of unique CpGs, gene expression probes and annotated genes present among the MEPs at each time point within the brackets.

13836, 83077, 15081 and 9 significant methylation-expression probe pairs (MEPs) (BH FDR P< 1x10^-4^) were identified comprising 10470, 63646, 10764 and 8 unique CpG-gene pairs at 1 week, 4 weeks, 8 weeks of exercise and 4 weeks of exercise withdrawal respectively. Of these 1535, 795, 677 and 8 independent genes and 82, 1827, 162 and 3 independent CpGs were present at 1 week, 4 weeks, 8 weeks of exercise and 4 weeks of exercise withdrawal respectively (Table 4). These represent 57%, 62%, 67% and 0.003% of DEGs (Fig 7A) and 0.07%, 3.6%, 0.09% and 0.003% of DMPs (Fig 7B) at exercise weeks 1, 4, 8 and exercise withdrawal respectively. This suggests DNA methylation may regulate a high proportion of gene expression changes in response to exercise at the concurrent timepoint, but via only a few CpGs across exercise timepoints, and gene expression changes post exercise withdrawal are primarily CpG methylation independent. There was no overlap between the MEPs across the time points (Fig 7C) or eQTM CpGs (Fig 7D), however, there was overlap between unique methylation associated genes at each time point (Fig 7E) suggesting that CpG specific association with gene expression is dynamic during exercise training. When considering the 1244 probes (779 genes) that were consistently differentially expressed across all time points (Fig 4A), 79.8% (993 gene probes, 675 genes) were associated with DNA methylation changes, implicating DNA methylation as an important mechanism regulating sustained changes in gene expression in response to exercise.

**Figure 7:**
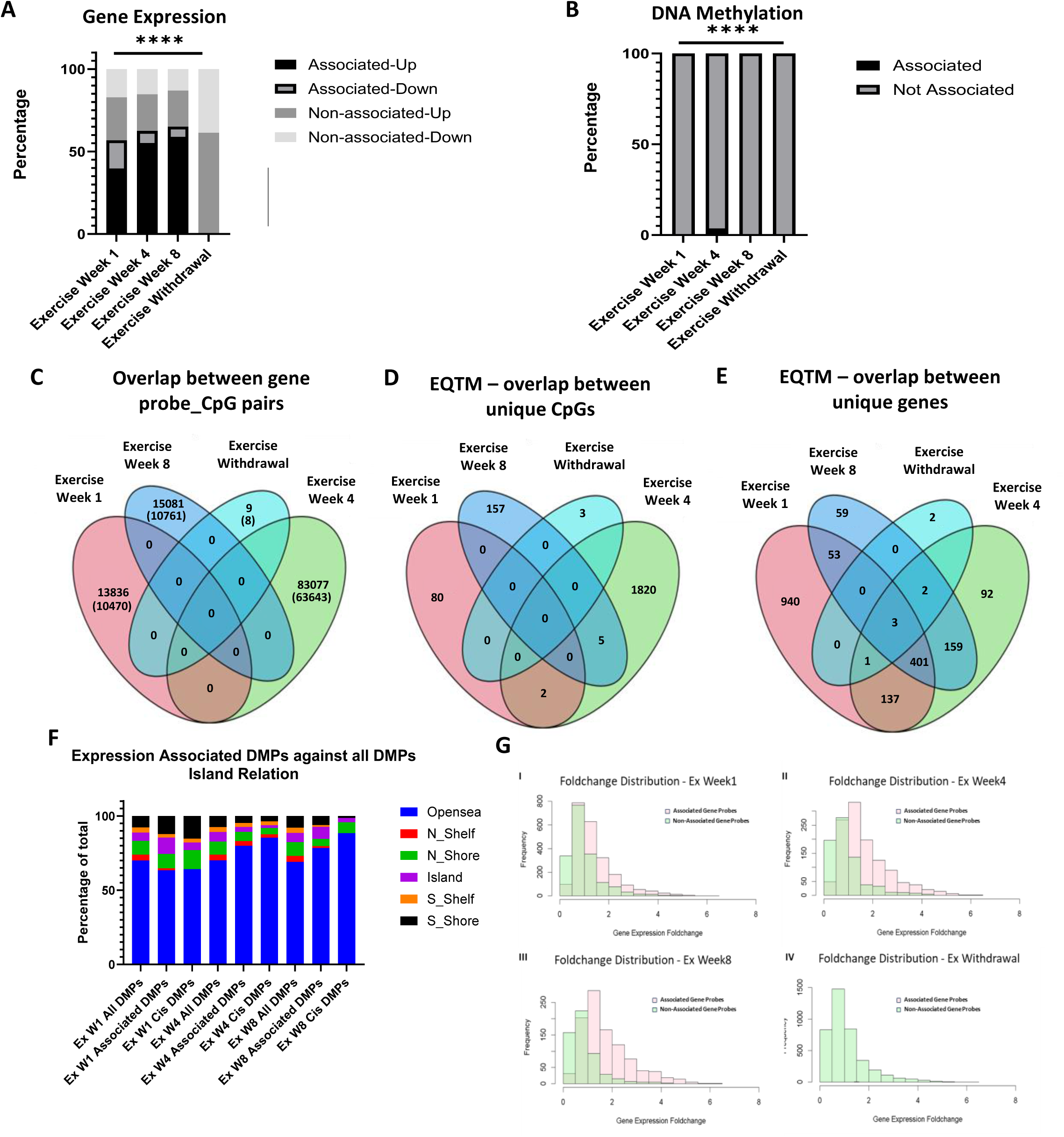
Characterisation of significant Methylation Expression Pairs (MEPs). **[A]** Contingency plot of methylation associated and non-associated DEGs representing percentage of genes that are either upregulated or downregulated showing significant difference in distribution across time points (****= Chi-Square test, p<0.0001). Methylation associated genes are predominantly upregulated. **[B]** Contingency plot depicting proportions of DNA expression associated and non-associated DMPs showing significant difference in distribution across timepoints (**** = Chi-Square test, p<0.0001) Venn Diagrams depicting **[C]** CpG-gene probe (number of unique genes depicted in brackets) pairs from eQTM analysis (Benjamini-Hochberg, p<1e-4), **[D]** unique expression associated differentially methylated CpGs across the time points and **[E]** unique methylation associated differentially expressed gene probes. **[F]** Contingency plot depicting distribution of CpG Island locations of expression associated differentially methylated. Chi-Square test revealed significant difference between the distribution of expression associated DMPs compared to all DMPs at week 4 (**** = p<0.0001).and week 8 (* = p<0.05). Expression associated DMPs at week 4 significantly enriched for opensea. **[G]** Absolute gene expression log foldchange distribution of DNA methylation associated (Pink) and non-associated (Green) gene probes for exercise week 1 (I), week 4 (II), week 8 (III) and exercise withdrawal (IV).

Upregulated genes were more likely to be DNA methylation associated than down regulated genes; 60%, 71%, 72% of upregulated genes at weeks 1,4 and 8 respectively versus 50%, 32% and 32% of the downregulated genes (Fig 7A). Both direct and inverse relationships were observed between CpG methylation and gene expression at exercise weeks 1, 4 and 8 (Fig EV2 A-C).

Gene expression associated CpGs significantly (Chi square p<0.0001) enriched for CpG islands at exercise week 1 and 8 (Fig 7F) suggesting that these methylation changes are located at gene regulatory regions. In terms of genomic location of CpGs relative to the associated gene, 5.2% (39 CpGs and 435 genes), 5.7% (592 CpGs and 502 genes) and 4.7% (73 CpGs and 293 genes) of the significant MEPs at exercise weeks 1, 4 and 8 respectively (Fig EV3 A-C, red points), and 1 out of the 9 MEPs identified at exercise withdrawal were present on the same chromosome. A greater number of methylation associated DEG probes exhibited larger gene expression effect size changes compared to non-associated counterparts at exercise weeks 1, 4 and 8 (Kolmogorov-Smirnoff test, p < 2.2e-16) (Fig 7G). These observations suggest that DNA methylation may be a key mediator of exercise associated gene expression regulation.

### DNA methylation-associated gene expression enriches for immunomodulatory and muscle tissue remodelling pathways

Subsequently, we aimed to understand the potential functional impact of gene expression changes specifically associated with DNA methylation, via pathway analysis of DNA methylation associated genes. DNA methylation associated genes were significantly enriched in 66, 52 and 35 pathways (p<0.05 and absolute(z-score)>1) at exercise weeks 1, 4 and 8 respectively (Appendix Table S7). 23 pathways enriched to methylation-associated genes at all exercise time points (Fig 8A). These included activation of inflammation and immune response pathways including *IL-4* signalling and macrophage alternative activation signalling (Fig 8B, green diamond); tissue remodelling pathways including integrin, *ILK*, actin cytoskeleton and *GP6* signalling; and fibrosis and tissue repair pathways including pulmonary fibrosis, hepatic fibrosis and wound healing signalling pathways (Fig 8B, purple diamond). In contrast, metabolism and oxidative stress response pathways and cytoskeleton/ cell motility and adhesion pathways were rarely associated with methylation changes at any time point.

**Figure 8:**
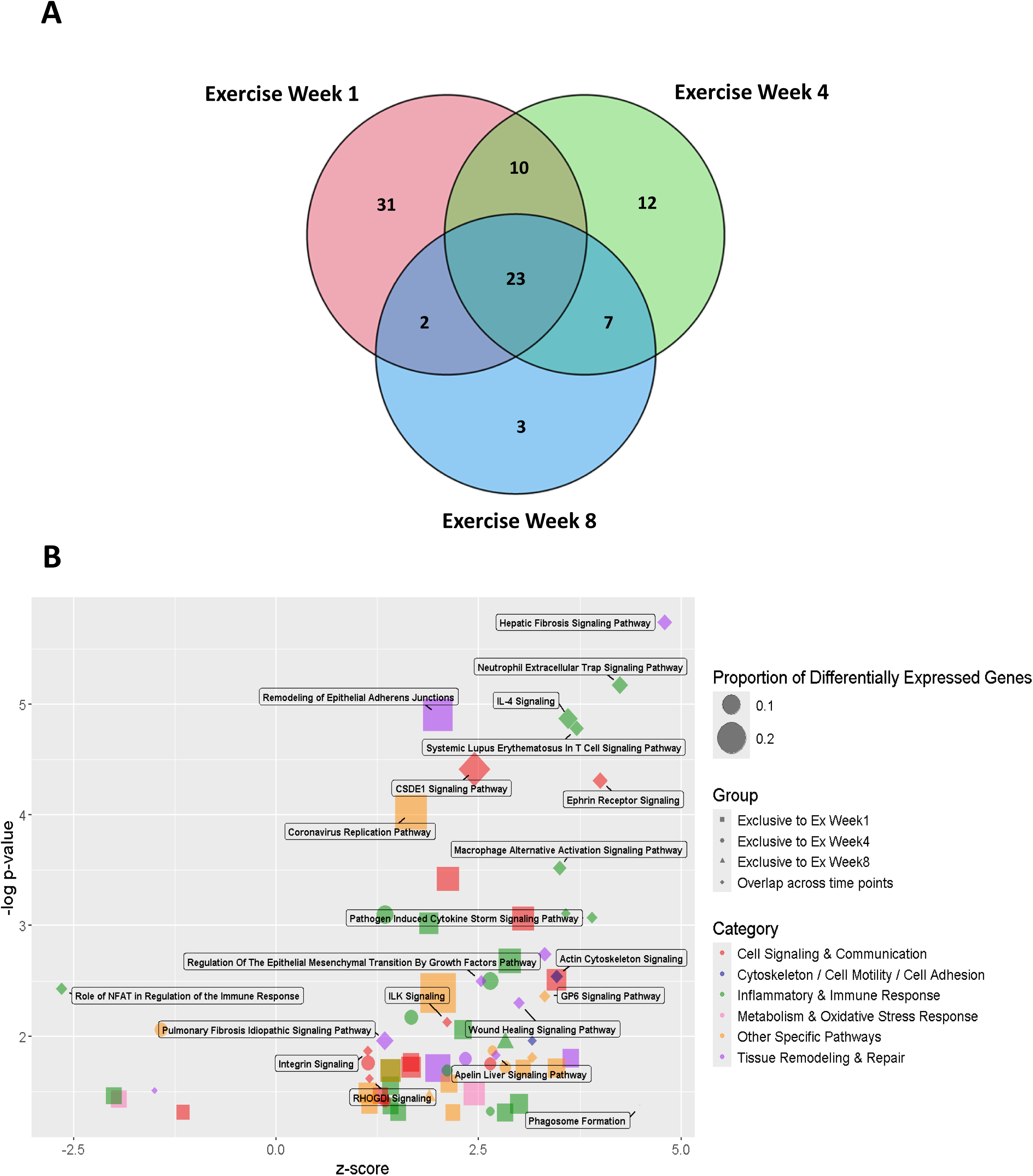
Pathway analysis of methylation associated differentially expressed genes performed using Qiagen Ingenuity Pathway Analysis software. Venn diagrams depicting **[A]** overlap of upstream regulators enriched to methylation associated DEGs at each time point; **[B]** A bubble plot of functional pathway associations for methylation associated genes that were exclusively enriched at each time point and those that were enriched across all time points, depicted by different shapes. The plot depicts −log(P value) and z-score on the axes, with the size of the bubble indicating the proportion of DEGs in the dataset comprised in the pathway.

The *IL-4* signalling pathway plays a crucial role in macrophage mediated muscle repair. The collagens (*COL12A1, COL13A1, COL1A1, COL1A2, COL3A1, COL4A1, COL4A2*, etc) (Kim *et al*, 2015; Robinson *et al*, 2017) and laminins (*LAMB1 and LAMA4*) (Damas *et al*, 2018; Hjorth *et al*, 2015) that were upregulated and enriched to *GP6*, wound healing and *IL-4* signalling pathways are upregulated post exercise in skeletal muscle (Kritikaki *et al*, 2021). This implicates DNA methylation as a molecular mechanism underlying muscle regeneration and repair pathways associated with exercise.

In addition, 985, 840 and 731 upstream regulators significantly enriched for exercise weeks 1, 4 and 8 respectively (Appendix Table S8) (Fig 9A). Fibrosis and exercise associated transcription factors Transforming growth factor beta (*TGFβ*), (Böhm *et al*, 2016; Han *et al*, 2021) and receptor tyrosine-protein kinase (*ERbB2*) (Amani *et al*, 2020) are among the top upstream regulators enriched for the DNA methylation-associated genes at all three time points (Fig 9B). Tumour suppressing transcription factor *THZ1* (Kuo *et al*, 2021) which is also an inhibitor of a myogenesis promoting transcription factor, Cyclin dependent kinase 7 (*CDK7*)(Ma *et al*, 2018) was predicted to be inhibited. In addition, exercise associated chemokine receptor 2 (*CCR2*) that modulates muscle damage response and elicits an essential immune response after an acute bout of exercise (Blanks *et al*, 2020) and DNA methylation regulated angiotensin-1 precursor AGT (Chaar *et al*, 2015; Demura *et al*, 2015) were upstream regulators enriched at later exercise time points of 4 and 8 weeks (Fig 9B).

**Figure 9:**
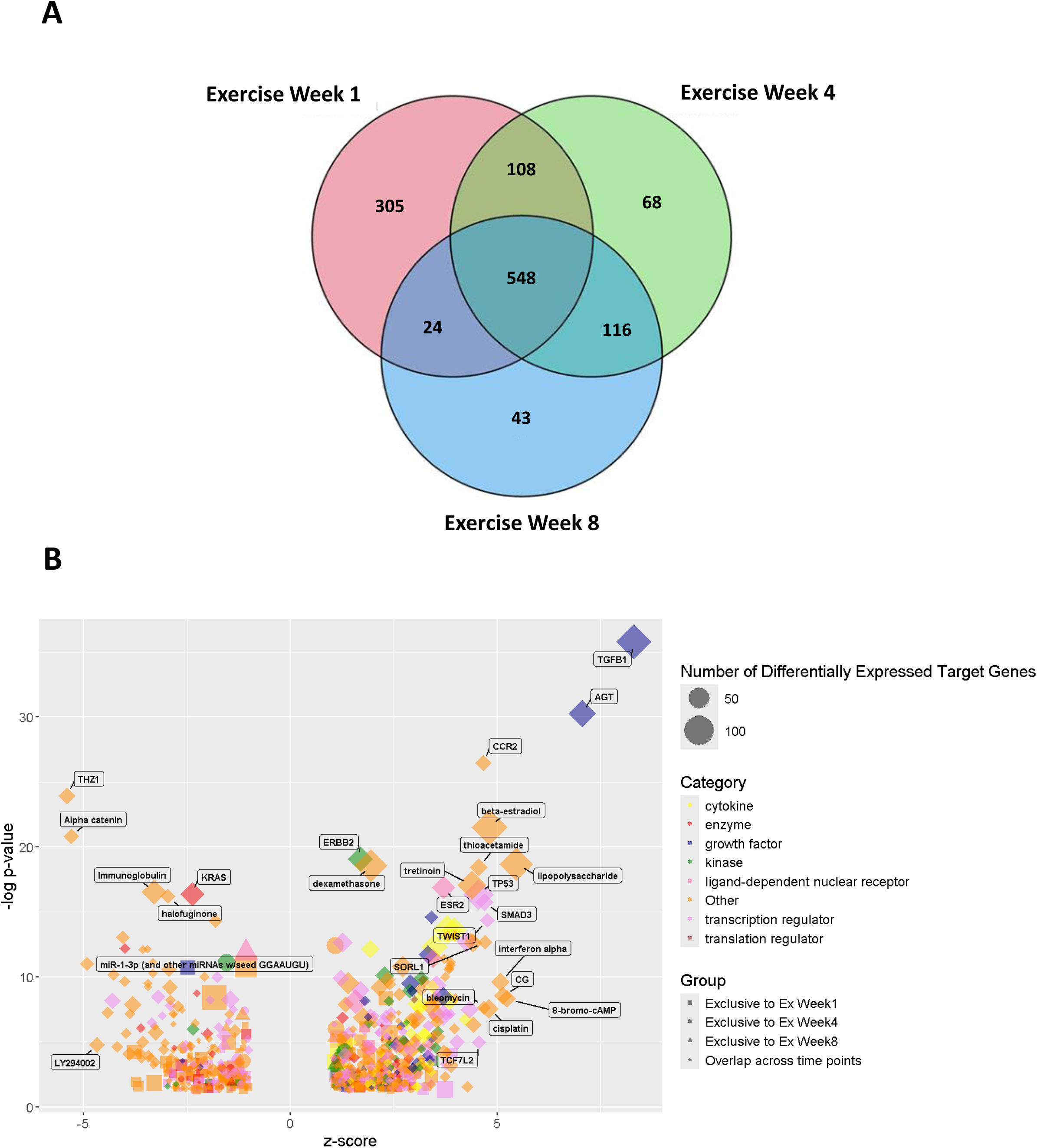
Upstream regulator enrichment analysis of methylation associated differentially expressed genes performed using Qiagen Ingenuity Pathway Analysis software. Venn diagrams depicting **[A]** overlap of pathways enriched to methylation associated DEGs at each time point; **[B]** A bubble plot of upstream regulator associations with methylation associated genes that were exclusively enriched at each time point and those that were enriched across all time points, depicted by different shapes. The plot depicts −log(P value) and z-score on the axes, with the size of the bubble indicating number of differentially expressed target genes in the dataset.

## Discussion

COPD patients exhibit impaired exercise tolerance and altered muscle metabolism, but the underlying molecular mechanisms, particularly in skeletal muscle, remain poorly understood. While DNA methylation changes have been associated independently with exercise, ageing and COPD across several tissues, its role in regulating skeletal muscle gene expression, in response to all three factors collectively has not been investigated in a comprehensive study. We examined transcriptome and DNA methylome changes in response to aerobic exercise training and withdrawal in, people with COPD sedentary healthy age matched and young volunteers to understand regulatory mechanisms of skeletal muscle response to exercise.

Here we demonstrate that while COPD status and age do not have significant impacts on skeletal muscle DNA methylation and gene expression profiles at baseline or in response to aerobic exercise training, exercise itself induces robust DNA methylation and gene expression changes. Transcriptomic changes that were transient, associated with acute pro-inflammatory and oxidative stress response pathways while DNA methylation-associated gene expression changes predominantly enriched for immunomodulatory and tissue remodelling/adaptation pathways that may contribute to changes in muscle function. While the DNA methylation independent transcriptomic changes reflect immediate responses to acute exercise that stabilise at later time points, the DNA methylation-associated gene regulation is linked to long-term adaptations of skeletal muscle in response to exercise. Overall, this study provides a deeper understanding of the molecular mechanisms underlying skeletal muscle response to a time course of aerobic training intervention, important for understanding how muscles adapt to training and utility in improving rehabilitation approaches in chronic disease management.

Our observation that there was no baseline differences in DNA methylation or gene expression between young healthy, COPD and age-matched older healthy individuals, is consistent with our previous report that intrinsic mitochondrial function is not significantly different between the groups (Latimer *et al*., 2022), suggesting skeletal muscle is more broadly (not just at the mitochondrial level) similar in aging and COPD, or that dysfunction (Jaitovich & Barreiro, 2018) arises via post-transcriptional mechanisms. In addition, the extent and clinical presentation of skeletal muscle dysfunction in COPD is heterogenous. Evidence indicates that muscle dysfunction can occur in early stages of the disease, while some individuals with severe COPD retain relatively preserved muscle function, highlighting a weak and non-linear relationship with lung function alone (Jaitovich & Barreiro, 2018). This variability is further driven by physical inactivity, systemic inflammation, ageing, and comorbidity profiles, thus explaining the complexity in unravelling the underlying mechanism (Vanfleteren *et al*, 2013). Age-associated DNA methylation changes have been reported in several tissues including blood, brain, liver, lung and kidney (Horvath, 2013; Horvath *et al*, 2012; Wang *et al*, 2023). A recent study investigated age-associated DNA methylation and associated gene expression changes in 9 tissue types including skeletal muscle and reported significant changes in all except skeletal muscle tissue (Jain *et al*, 2024). However, other studies have shown decreased methylation in aged skeletal muscle tissue (Turner *et al*., 2020; Zykovich *et al*., 2014) and developed a dedicated epigenetic clock for skeletal muscle (Voisin *et al*, 2020). In our study, the lack of significant differences in DNA methylome and transcriptome between young and older healthy individuals was reported at our genome wide significance threshold of 9x10^-8^. At a lower significance of Benjamini-Hochberg FDR<0.05 41,297 DMPs (Appendix Fig S1A) and 416 DEGs (Appendix Fig S1B) were present, indicating a potential age effect below our statistical rigour, as such we included age as a covariate in all our exercise effect analyses.

We have demonstrated that COPD or age does not influence transcriptome and methylome changes in response to exercise in skeletal muscle at any point during eight weeks of exercise training and following 4 weeks of exercise withdrawal. This is consistent with our previous targeted gene expression profiling of 59 transcripts involved in muscle fuel metabolism, that showed no age or disease specific differences in expression in response to exercise, despite functional temporal differences in intrinsic mitochondrial muscle function, in the three groups (Latimer *et al*., 2022), suggesting post-transcriptional mechanisms may be responsible for the age and COPD associated alteration to mitochondrial function. This observation is also supported by findings from a controlled bed rest immobilization study in healthy men, which showed that early and pronounced changes in skeletal muscle mRNA expression during disuse were not uniformly mirrored by corresponding physiological and metabolic adaptations, highlighting that early gene expression changes to an intervention are not always directly concordant with downstream or subsequent functional outcomes (Shur *et al*, 2022). A limitation of the present study is the lack of proteomic analyses to directly assess post-transcriptional regulation. The study of Latimer er al (Latimer *et al*., 2022) also described differences in whole body responses (V’_O2peak_) in healthy individuals versus those with COPD. Here we focus on muscle gene expression and DNA methylation which are several measures of function away from being applicable to whole body responses, preventing us from drawing conclusions in this context. However, accelerated epigenetic and transcriptomic ageing does have a bidirectional relationship with diseases. Specifically, epigenetic age acceleration was observed in skeletal muscle in individuals with chronic liver disease (Nicholson *et al*, 2024) and another study demonstrated that accelerated epigenetic ageing in peripheral blood leukocytes correlated with increase cardiovascular risk factors (Ammous *et al*, 2021). Exercise is known to be a cost effective and efficient intervention to alleviate ageing associated morbidities and promotes healthy ageing (Cartee *et al*, 2016). Evidence suggests that baseline aerobic fitness correlates with younger epigenetic and transcriptome age, with aging-associated changes reversible by exercise (Liang *et al*, 2024; Voisin *et al*, 2024). Specifically in COPD individuals, incorporating supervised exercise in pulmonary rehabilitation improves overall health-related quality of life (Costes *et al*., 2015; Pothirat *et al*., 2015; Ward *et al*, 2025). In this study, it is important to note that some individuals with COPD had lower muscle loading based on their baseline ventilatory limitations. The exercise training intervention was individually prescribed and supervised resulting in measurable whole-body physiological adaptation in all groups (Latimer et al, 2022).

Subsequently we investigated the integrated response of methylome and transcriptome on aerobic exercise irrespective of COPD and age through the course of 8 weeks of exercise and following detraining. Aerobic, endurance and resistance exercises are known to induce transcriptomic alterations in skeletal muscle which progressively shapes muscle function as well as overall health benefits and protection against chronic diseases. Some recent studies are consistent with our findings of transcriptomic changes associating with metabolism and oxidative stress response (Egan & Sharples, 2023; Pillon *et al*, 2020), immunomodulation (Rubenstein *et al*, 2022) and tissue remodelling (Zeng *et al*, 2023) functions post aerobic training. Specifically, acute aerobic exercise training is associated with mitochondrial respiration, lipid metabolism, glucose transport and antioxidant functions (Egan & Sharples, 2023). This is consistent with our findings after 1 week of exercise. Another study reported similar changes in inflammatory response to acute exercise in skeletal muscle of active and sedentary older adults, consistent with our findings (Rubenstein *et al*., 2022). In addition, we observed stable changes over the period of 8 weeks that persisted following exercise cessation for 4 weeks, predominantly associated with immunomodulatory and tissue remodelling pathways. A comparative study of acute versus chronic aerobic training revealed that chronic training elicited more pronounced transcriptomic alterations, particularly in immunomodulatory and extracellular matrix remodelling pathways, among older adults (Zeng *et al*., 2023). Accompanying these robust transcriptomic changes, transient DNA methylation changes were observed after exercise week 1 and more persistent changes appeared at later time points.

Association between DNA methylation and gene expression is complex and previous studies have investigated targeted DNA methylation promoter sites or sites annotated to exercise associated DEGs (Alibegovic *et al*., 2010; Nitert *et al*., 2012; Turner *et al*., 2019). We chose to use an expression quantitative trait methylation approach, previously unreported in exercise and skeletal muscle associated ‘omics studies as tests for associations between each DEG and DMP identified, generating a complete analysis of gene-methylation interactions without restricting to overlap in gene level annotations in transcriptome and methylome changes. Using this method, we identified genes that are associated with methylation changes suggesting regulation by DNA methylation. Heart Development Protein with EGF like Domains 1 gene (*HEG1*), a blood vessel and heart development (Kleaveland *et al*, 2009) and malignant cell growth regulating gene (Tsuji *et al*, 2017), was observed in our data as one of the consistent DEGs that persisted even after exercise withdrawal. In addition, it also significantly associated with distinct DMPs at exercise weeks 1 (3 DMPs), 4 (11 DMPs) and 8 (4 DMPs). A time course study in young healthy males who underwent acute and chronic resistance exercise training followed by detraining and retraining demonstrated a phenomenon of epigenetic memory, some of which linked to gene expression changes (Seaborne *et al*., 2018). HEG1 was reported as one of the epigenetic memory genes, thus suggesting our observation of distinct methylation associations across time points as a complex cascade of epigenetic regulatory mechanism that maintains the expression of HEG1 even after exercise withdrawal. Although previously unreported in relation to muscle function, the role of this gene as an epigenetic memory mark induced in both resistance (Seaborne *et al*., 2018) and aerobic training exercises encourages further investigation. With this as an exemplar gene that showed persistent methylation associated expression change, we have reported 779 genes that were consistently differentially expressed across all time points and were predominantly associated with differential DNA methylation. These genes were primarily associated with inflammation, immunomodulatory and muscle tissue remodelling pathways. Several studies have reported ECM adaptation and muscle remodelling post exercise. Specifically, a recent study reported alterations in intramuscular ECM protein expression in individuals with COPD in response to exercise training compared to healthy adults, suggesting that impaired ECM remodelling may underlie the poor muscle adaptation observed in COPD (Simoes *et al*, 2025). Another study investigated the transcriptome changes after 12 weeks of endurance and strength training in middle-aged sedentary males and reported 28 genes that directly related to ECM remodelling including the genes enriching to *GP6* and *IL-4* pathways reported in our study (Hjorth *et al*., 2015). It has also been reported that running altered gene expression of ECM modulating collagens and laminins in cortex and hippocampus of young but not mid-life mice (Foley *et al*, 2019). This overlaps with our findings and the specific laminin and collagen genes enriched to *GP6* and hepatic fibrosis signalling pathways. Further understanding of these pathways, constituent genes, and regulation by DNA methylation in relation to exercise response in skeletal muscle will unravel new insights into the mechanisms of muscle remodelling post exercise. DNA methylation associated and independent transcriptomic changes associated with tissue remodelling and muscle adaptation may differentially influence cellular and physiological responses in healthy and COPD individuals; however, further studies are required to elucidate the intermediate steps linking transcriptional changes to cellular and physiological function. Overall, this study offers more molecular granularity compared to other existing studies that focussed on acute exercise effect specifically acting as a homeostasis perturbation stimulus that triggers a cascade of immune response and oxidative stress response pathways (Laker *et al*., 2017; Turner *et al*., 2019). It provides increased insight into the molecular basis of aerobic exercise response across the time course of an eight-week training programme in skeletal muscle, highlighting a shift from transient pro-inflammatory and oxidative stress response early in the training programme to a sustained immunomodulatory and tissue adaptative transcriptomic change that is potentially regulated by DNA methylation.

## Methods

### Demographics

The study cohort comprised of 9 young healthy volunteers, 20 COPD and 10 age matched older healthy volunteers. Full demographics are provided in Table 1. All individuals had a sedentary lifestyle devoid of pulmonary rehabilitation or exercise training 1 year prior to the study. Habitual step-counts were matched in all participants. Testing and training was conducted at Glenfield Hospital, Leicester and University of Nottingham Medical School, QMC site, Nottingham. Participants gave written informed consent, and the study was approved by NHS Research Ethics Committee West Midlands –Coventry & Warwickshire, Reference 13/WM/0075. Full inclusion criteria are previously published (Latimer *et al*., 2022) and the demographics relevant to the current study are detailed in Table 1.

### Exercise Training

A non-randomised interventional longitudinal cohort study was performed as described previously (Latimer *et al*., 2022). In summary, all participants performed supervised cycling exercise on a stationary cycle ergometer (Lode, Groningen, Netherlands) three times a week for a period of eight weeks. Supervised training was withdrawn after eight weeks and participants were instructed to resume their pre-study sedentary lifestyle for a period of 12 weeks (Figure 1).

### Skeletal Muscle Biopsy

A microbiopsy of vastus lateralis muscle was taken 24 hours post supervised exercise training at rested and fasted state. Biopsies were collected at baseline (before starting the exercise training); after 1, 4 and 8 weeks of supervised exercise training; and 4 weeks after subsequent exercise withdrawal. Biopsy tissue was carefully subject to microdissection to remove any non-muscle tissue, and processed for DNA and RNA extraction

### DNA and RNA Isolation

DNA was isolated using Qiagen QIAamp DNA mini kit according to the protocol provided by the manufacturer and previously described (Constantin-Teodosiu *et al*, 2020). RNA was isolated from snap frozen tissue using TRI reagent (Ambion, Applied Biosystems) according to the manufacturer’s protocol. Briefly, the tissue was homogenised in 1ml of TRI Reagent and 20ul of 10ug/ul glycogen solution, phase-separated with 200ul of chloroform, and the aqueous phase was precipitated with 500ul isopropanol. The RNA pellet was washed with 75% ethanol, air-dried, and resuspended in 50ul of RNase-free water.

### Bisulfite conversion and DNA methylation arrays

Genomic DNA (750ng) was subject to bisulfite conversion using the EZ DNA Methylation Kit (Zymo Research) as per the manufacturer’s instructions. Specific incubation conditions for the Illumina Infinium Methylation Assay were applied as per the manufacturer’s protocol Appendix. Samples were eluted in 12 μl of the elution buffer. Bisulfite-converted DNA wasquantified using a NanoDrop^TM^ 8000 Spectrophotometer (Thermo Fisher Scientific). 160 ng of the conversion product was used for DNA methylation quantification at over 850,000 CpG sites using the Illumina Infinium HumanMethylationEPIC BeadChip array, according to the manufacturer’s protocols.

### DNA methylation data quality control and normalisation

Arrays run on the HumanMethylationEPIC BeadChip were scanned using an Illumina HiScan scanner. Raw IDAT files obtained through GenomeStudio software were imported into R Statistical software ((2020)) using minfi package (version 1.52.1) (Aryee *et al*, 2014; Fortin *et al*, 2017). Pre-processing and quality control steps were carried out using minfi package as follows. Preliminary quality control steps identified no samples that required exclusion based on low median intensities, sex mismatch or distinct outliers in the beta distribution plot. Technical replicates of eight samples displayed robust correlation (0.994 ≤ r ≤ 0.997). Functional between-sample normalisation was performed using the *preprocessFunnorm (Fortin et al, 2014)* function in minfi package. Correlation of technical replicates improved post normalisation (0.996 ≤ r ≤ 0.997). 27,889 probes with less than three bead count in at least one sample were removed. Subsequently, further probe filtering removed probes comprising 65 single nucleotide polymorphism probes (SNP), 2,180 probes with low detection p-value (0.05) in at least one sample, 18,537 probes that were mapped to sex chromosomes, 11,512 probes that bind to autosomal and sex chromosomes, 283 probes with a SNP at a targeted CpG binding sites, 39,577 cross-hybridising probes and 2464 non-CpG targeting probes (Pidsley *et al*, 2016). 766,569 probes remained for analysis. Sequential batch effect correction based on array chip ID and sample position on the chip was performed using using *comBat* function in SVA package (version 3.54.0) (Leek *et al*, 2012) and correlation of beta values before and after batch correction was >0.997 in all samples. Two sets of DNA methylation measures were calculated – Beta values (ranges between 0 to 1 and represents the ratio of methylated signal to total signal intensity) representing percentage methylation and M values (log transformed beta values) which are robust measurements used for statistical analysis.

### Gene Expression Microarray

Whole-genome transcriptome analysis was conducted by GlaxoSmithKline using the Affymetrix Human Gene U133 Plus 2.0 Array Strips (Affymetrix, Santa Clara, CA, USA). Raw CEL files were imported into R statistical software using Affy package (version 1.84.0) (Gautier *et al*, 2004) and subject to GC Robust Multiarray Average (GCRMA) normalisation using gcrma package(version 2.78.0) (Gentry, 2020). Following quality control checks after normalisation, principal component analysis revealed two non-specific clusters that did not associate with any covariates which were adjusted for using the *comBat* function in SVA package (version 3.54.0) (Leek *et al*., 2012).

### Differential Analysis

Differential CpG methylation and gene expression were identified by multi-level linear regression analysis (empirical bayes method) including within subject correlation for matched samples across time points using the limma package (version 3.62.1) (Ritchie *et al*, 2015) in R. Smoking status was used as covariate in all linear regression models. Five separate analyses were performed for DNA methylation and gene expression as described in Table 2 and 3 respectively. A Benjamini-hochberg false discovery rate (BH FDR) less than 9x10^-8^ was taken as significant (Mansell *et al*, 2019).

### Integration of Transcriptome and Methylome Data

Association between DEGs and differentially methylated CpGs was established using Expression Quantitative Trait Methylation analysis (eQTM), using simple linear regression in the limma package (version 3.62.1) without any covariates. The gene-CPG pairs with BH FDR <1x10^-4^ were considered significant methylation-expression associations (Methylation-Expression pairs – MEPs).

### Pathway Analysis

Pathway analysis was performed on DEGs and methylation associated genes using Qiagen Ingenuity Pathway Analysis software. A z-score cut off 1 was applied.

## Data availability

The datasets and computer code will be available before publication

## Acknowledgements

The authors gratefully acknowledge the contribution to this work of: The human studies technical staff in the David Greenfield Human Physiology Unit, University of Nottingham; Glenn Hearson, NIHR Nottingham BRC respiratory theme; all participants who volunteered for this study; and Bruce Miller, Ruth Tal-Singer, Divya Mohan, Ashutosh Pandey and Michal Magid-Slav for generating the gene expression data.

## EXPANDED VIEW

**Figure EV1:**
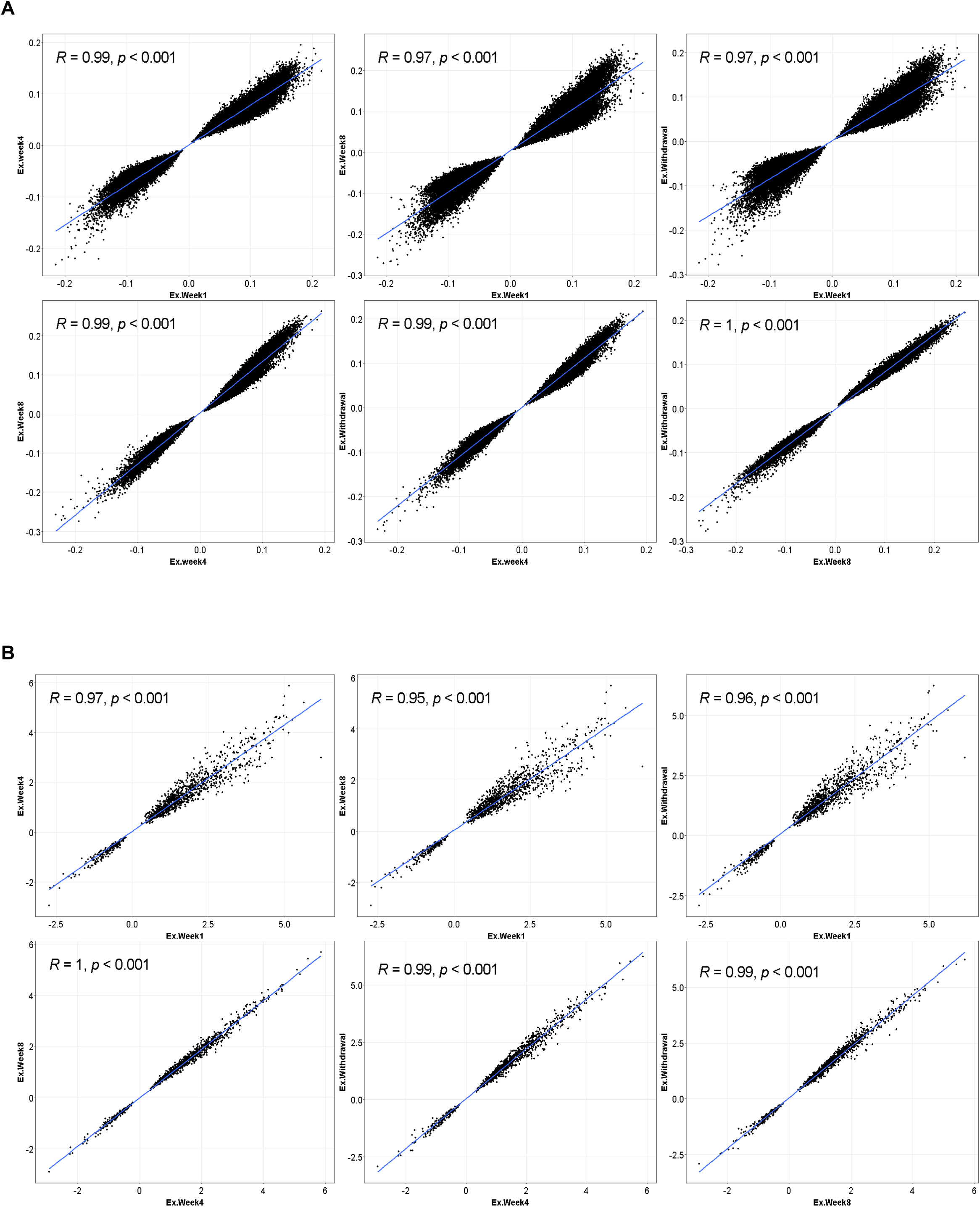
39112 differentially methylated CpG probes and 1244 gene probes overlapped between all exercise time points. **[A]** Correlation of deltabeta (corresponding to baseline) values of these overlapped CpGs between each pair of exercise time points show consistent change in direction with significant Pearson correlation greater than 0.97 (p<0.001). **[B]** Correlation of gene expression foldchange (corresponding to baseline) values of these overlapped gene probes between each pair of exercise time points show consistent change in direction with significant Pearson correlation greater than 0.95 (p<0.001).

**Figure EV2:**
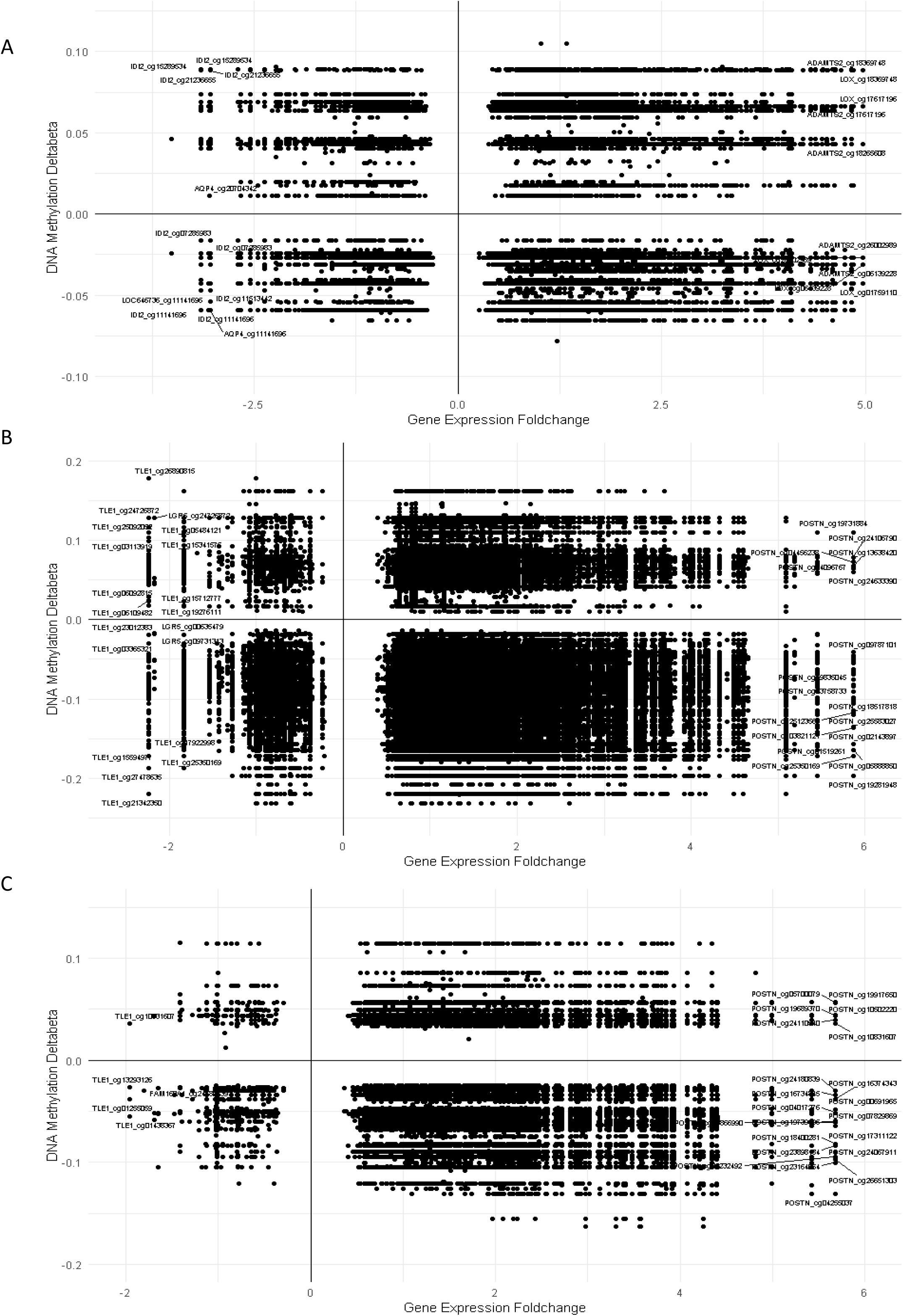
Investigating the directional relationship between methylation and expression associations. Plot of paired gene expression foldchange and DNA methylation beta values depicting the direct or inverse relationship between gene-cpg pairs from eQTM analysis for each time point – Exercise Week 1 **[A]**, Exercise Week 4 **[B]** and Exercise Week 8 **[C]**.

**Figure EV3:**
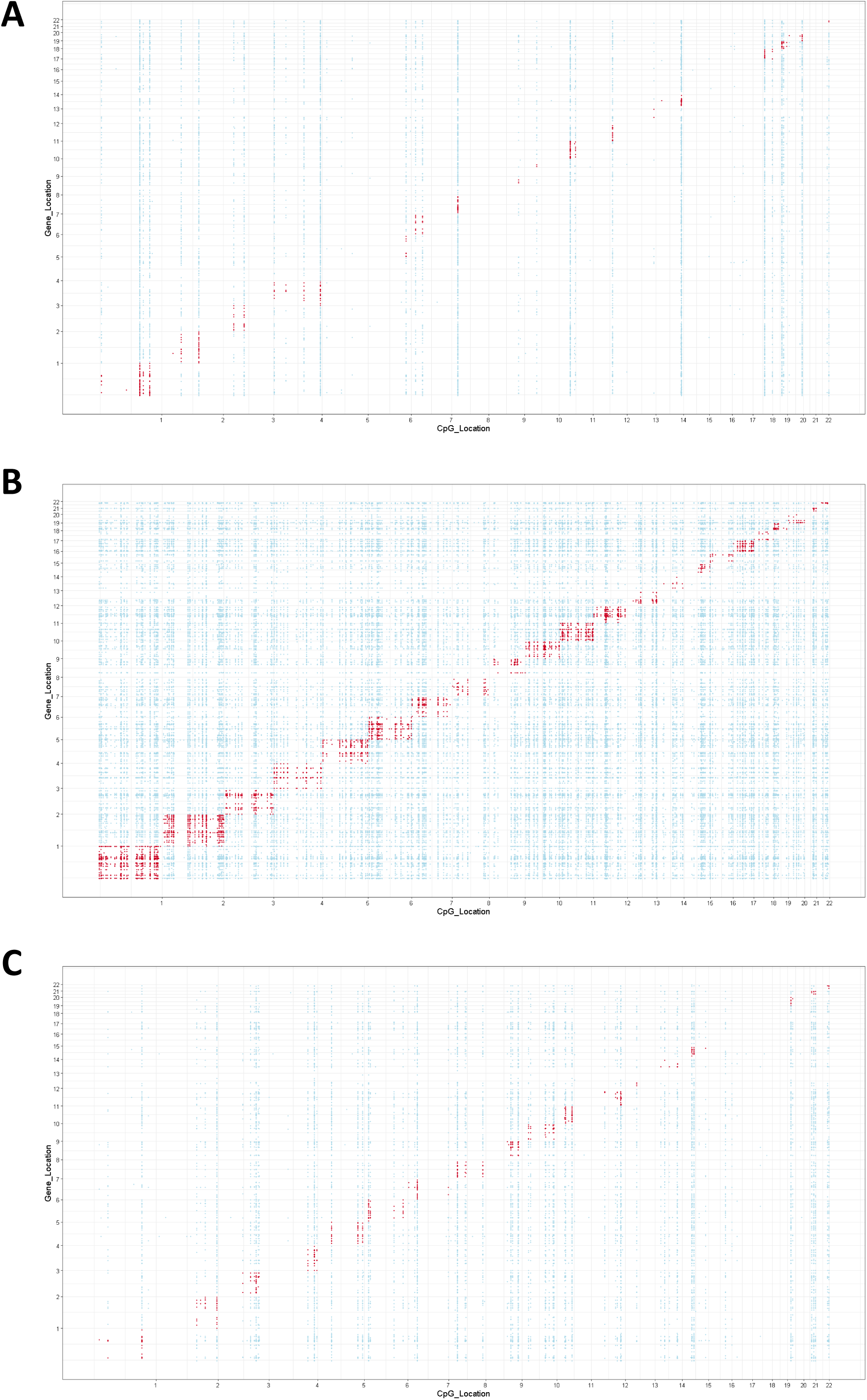
Mapping genomic locations of methylation and gene expression associations. Plot of CpG genomic location and transcription start site location of MEPs at exercise week 1 **[A]** exercise week 4 **[B]** and exercise week 8 **[C].** Trans associations defined as gene and CpG pairs located on different chromosomes are depicted as blue plot points and cis association defined as gene and CpG pairs located on same chromosomes are depicted as red plot points.

## APPENDIX

**Appendix Table S1 :**
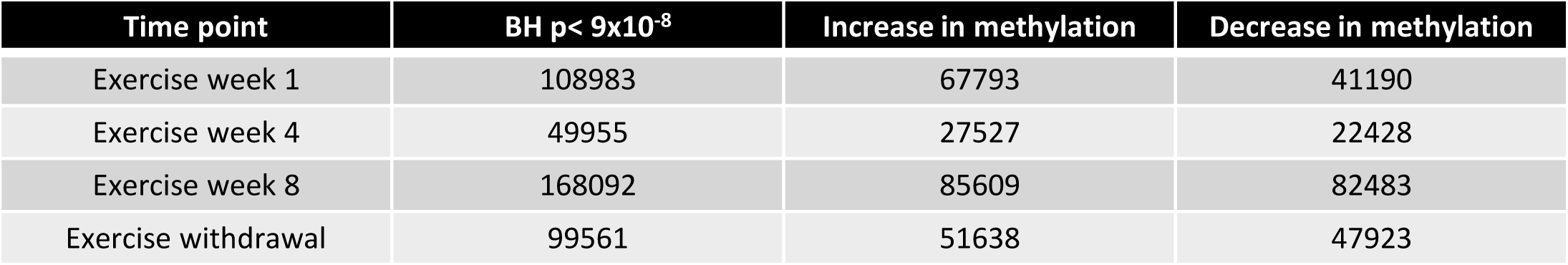
Table of results for multi-level linear modelling investigating the effect of exercise on DNA methylation of skeletal muscle. The table depicts the number of CpGs that were differentially methylated and number of significant DMPs that showed an increase/decrease in methylation compared to baseline for each time point. Sites were taken as significant at Benjamini-Hochberg FDR <9e-8

**Appendix Table S2 :** Table of results for multi-level linear modelling investigating the effect of exercise on gene expression of skeletal muscle. The table depicts the number of genes that were differentially expressed, number of annotated genes among the significant DEGs and number of significant DEGs that showed an increase/decrease in expression compared to baseline for each time point. Probes were taken as significant at Benjamini-Hochberg FDR <9e-8

| Time point | BH $p < 9 \times 10^{-8}$ | Increase in expression | Decrease in expression |
| --- | --- | --- | --- |
| Exercise week 1 | 4413 (annotated: 3968) | 1357 | 2611 |
| Exercise week 4 | 2081 (annotated: 1898) | 1468 | 430 |
| Exercise week 8 | 1682 (annotated: 1537) | 1242 | 295 |
| Exercise withdrawal | 4406 (annotated: 3972) | 2435 | 1537 |

**Appendix Figure S1:**
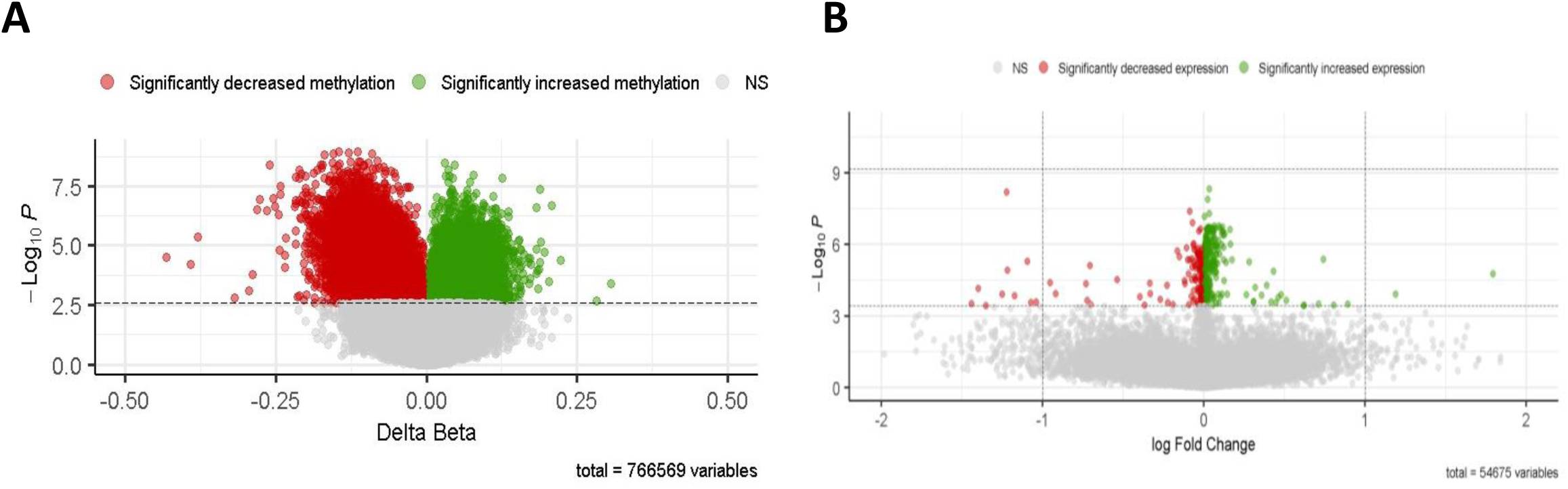
Effect of age on skeletal muscle DNA methylation, gene expression and epigenetic age. **[A]** Plot of 766569 DNA methylation probes used in the analysis depicting 41297 significantly differentially methylated sites (Benjamini-Hochberg FDR, p<0.05) in response to age at baseline. **[B]** Plot of 54675 gene expression probes used in the analysis depicting 416 significantly differentially expressed genes (Benjamini-Hochberg FDR, p<0.05) in response to age at baseline. Red/Green = Benjamini-Hochberg FDR < 0.05. The horizontal dotted lines indicate the p value corresponding to Benjamini-Hochberg FDR < 0.05. Red = decreased methylation/expression at exercise timepoint, green = increased methylation/expression at exercise timepoint.

